# Unique belowground ecological strategies of subtropical and tropical plant species expand the root trait space

**DOI:** 10.1101/2024.10.06.616893

**Authors:** Nathaly R. Guerrero-Ramírez, Monique Weemstra, Shalom D. Addo-Danso, Kelly Andersen, Marie Arnaud, Amanda L. Cordeiro, Daniela F. Cusack, Martyna M. Kotowska, Ming Yang Lee, Céline Leroy, Laynara F. Lugli, Kerstin Pierick, Chris M. Smith-Martin, Laura Toro, María Natalia Umaña, Oscar J Valverde-Barrantes, Michelle Wong, Claire Fortunel

**Affiliations:** Biodiversity, Macroecology and Biogeography, Faculty of Forest Sciences and Forest Ecology, University of Göttingen, Göttingen 37077, Germany; Silviculture and Forest Ecology of Temperate Zones, Faculty of Forest Sciences and Forest Ecology, University of Göttingen, Göttingen 37077, Germany; Centre of Biodiversity and Sustainable Land Use, University of Göttingen, Göttingen 37077, Germany; Department of Biological Sciences, International Center for Tropical Biodiversity, Florida International University, Miami, FL 33199, USA; Department of Ecology and Evolutionary Biology, University of Michigan, Ann Arbor, Michigan 48103, USA; Forests and Climate Change Division, CSIR-Forestry Research Institute of Ghana. P.O Box UP 63, KNUST, Kumasi, Ghana; College of Science, Nanyang Technological University, Singapore, Singapore; Institute of Ecology and Environmental Sciences Paris (iEES-Paris), Sorbonne University, Paris, France; Birmingham Institute of Forest Research & School of Geography, Earth & Environmental Sciences, University of Birmingham, Birmingham, UK; Department of Ecosystem Science and Sustainability, Colorado State University, CO, 80523, USA; Smithsonian Tropical Research Institute, Balboa, Ancon 0843-03092, Republic of Panama; Department of Plant Ecology and Ecosystem Research, University of Göttingen, Göttingen, Germany; School of Natural Sciences, Macquarie University, Sydney, Australia; Asian School of the Environment, Nanyang Technological University, Singapore; AMAP, Université de Montpellier, CIRAD, CNRS, INRAE, IRD, France; EcoFoG, AgroParisTech, CIRAD, CNRS, INRAE, Université des Antilles, Université de Guyane, Campus agronomique, Kourou, France; Land Surface-Atmosphere Interactions, Technical University of Munich, Hans-Carl-von- Carlowitz-Platz 2, 85354, Freising, Germany; Spatial Structures and Digitization of Forests, Faculty of Forest Sciences and Forest Ecology, University of Göttingen, Göttingen 37077, Germany; Department of Plant and Microbial Biology, University of Minnesota, St. Paul, Minnesota, 55108, USA; Missouri Botanical Garden, St. Louis, Missouri, 63110, USA; Department of Ecology and Evolutionary Biology, Yale University, New Haven, CT 06511, U.S.A; Cary Institute of Ecosystem Studies, Millbrook, NY 12545, U.S.A; AMAP (Botanique et Modélisation de l’Architecture des Plantes et des Végétations), Université de Montpellier, CIRAD, CNRS, INRAE, IRD, Montpellier, France

**Keywords:** biogeography, functional diversity, phylogeny, root economics space, root functional traits, subtropics, tropics

## Abstract

Root trait variation may reflect the ecological and evolutionary processes shaping biodiversity, but remains poorly quantified in the (sub)tropics. Here, we aim to further complete our knowledge of belowground functional strategies by assessing the contributions of subtropical and tropical species to global root trait diversity. We gathered root data for 1618 temperate, 341 subtropical, and 775 tropical species. We compared functional diversity among biomes and calculated the unique contribution of each biome to the global root economics space. Further, we determined if the within-variation of subtropical and tropical biomes is shaped by species’ niches and/or differences in evolutionary history. Root trait expressions differed among biomes, but root functional diversity did not. Furthermore, subtropical and tropical biomes accounted for 40% of the unique root functional space within the global traits space. Species’ climate niches and phylogenetic turnover explained variation in root traits (e.g., denser root tissue was associated with drier sites) among subtropical but not tropical species. Through their unique root traits, sub(tropical) species strongly expand the current ‘global’ root trait space. This work underwrites their importance in conceptual models for more complete insights into how various belowground strategies drive plant functional biogeography and biodiversity globally.

## Introduction

Subtropical and tropical biomes are hotspots of biodiversity across taxonomic groups. Recent estimates suggest that tropical regions in America, Africa, and Asia jointly harbor ∼224,000 species amounting to ∼65 % of the known terrestrial vascular plant species (Raven et al. 2020, Govaerts et al. 2021). The large contribution of tropical ecosystems to global biodiversity may be associated with the high functional diversity in these systems, owing to *e.g.,* biogeographical transition zones in the subtropics where species with temperate and tropical strategies can coexist (Morrone 2024); their long evolutionary time and strong competition and niche partitioning; and their abundance of rare species with distinct plant ecological strategies that may minimize niche overlap, enabling the coexistence of a large number of functionally distinct species (Leitão et al. 2016, Umaña et al. 2017). Trait diversification may be more pronounced belowground than aboveground because root traits are more flexible in their coordination, giving rise to great variability in root trait combinations compared to leaf traits (Kramer-Walter et al. 2016). As data on root traits lag behind data on leaves in general but especially in the tropics (Cusack et al. 2024), our current insights on functional diversity and how it underlies species diversity and coexistence in (sub)tropical biomes may only represent the tip of the iceberg and remain largely incomplete. Here, we explore the emergent root trait variation among species across subtropical and tropical biomes from around the globe, how much it contributes to global belowground functional diversity, and to what extent it is driven by biogeographic and evolutionary histories at the continental scale.

Interspecific variation in functional trait expressions generally reflects species’ variation in ecological strategies (Grime 1977). Ecological frameworks approach this relationship from an ‘economics’ perspective based on resource returns on carbon (C) investments into plant tissue (Bloom et al. 1985). Species invest either in the construction of robust, durable tissue with slow rates of resource return or in producing short-lived tissue with fast resource returns on relatively low biomass investment (Reich et al. 1997, Westoby et al. 2002, Wright et al. 2004). This tradeoff is evidenced by the global ‘leaf economics spectrum’ that spans a continuum of species that exhibit a suite of ‘slow’ versus ‘fast’ leaf traits (Reich et al. 1997, Wright et al. 2004) and is associated with high survival rates in adverse environments versus high growth rates on productive sites, respectively (Reich et al. 1998, Aerts and Chapin III 2000, Poorter and Bongers 2006). The traits of absorptive roots (i.e., fine roots responsible for nutrient and water acquisition and therefore, functional analogs of leaves; hereafter: roots) have been assumed to be similarly coordinated along a fast-slow axis in a ‘root economics spectrum’ (Mommer and Weemstra 2012, Reich 2014), but more recent work shows that root trait covariation is multidimensional (Kramer-Walter et al. 2016, Weemstra et al. 2016, Bergmann et al. 2020). A global analysis of root trait data showed that interspecific covariation in root traits was best captured by two independent axes in a ‘root economics space’ (Bergmann et al. 2020). The ‘collaboration axis’ is assumed to reflect the extent to which species rely on (arbuscular) mycorrhizal symbiosis for nutrient and water acquisition. It ranges from species with a high specific root length (SRL, root length per unit root dry mass) that enhances direct water and nutrient uptake by roots (*i.e.*, the do-it-yourself strategy), to species with thick roots that rely more heavily on mutualistic associations for resource capture (*i.e.*, the outsourcing strategy). The ‘conservation axis’ resembles the traditional fast-slow tradeoff, separating species that have tough, long-lived roots (*e.g.*, by having high root tissue density, RTD) with slow resource returns on C investments from those with resource-acquisitive roots with low biosynthesis costs and fast returns on C investments (*e.g.*, by having high N concentrations and thus, assumingly, fast metabolism) (Bergmann et al. 2020). This global multidimensional belowground framework is increasingly serving as a baseline for comparative approaches in functional ecology.

The root economics space was tested using 748 woody and non-woody species sampled around the globe but came with the major caveat that tropical and subtropical species are vastly underrepresented (Bergmann et al. 2020). As such, this lack of data and knowledge on (sub)tropical root trait (co-)variation hinders our understanding of overall ecological processes and mechanisms (Freschet et al. 2017, Bergmann et al. 2020, Carmona et al. 2021). Taking this bias towards the temperate biome into account is particularly relevant since (co)variation in root traits may well deviate among biomes (Gu et al. 2014, Weemstra et al. 2023) owing to different environmental conditions and unique evolutionary processes that select for different trait expressions (Comas et al. 2012, Eiserhardt et al. 2017, Valverde-Barrantes et al. 2021). Biotic and abiotic differences among biomes, such as nitrogen (N) versus phosphorus (P) limitation or co-limitation (Vitousek 2004), nutrient cycling rates (Marklein et al. 2016), mycorrhizal associations for the dominant canopy species (Steidinger et al. 2019, Soudzilovskaia et al. 2019), and the length of the growing season suggests that trait syndromes found in temperate areas may not represent belowground functional diversity in lower latitudes.

Improved data availability already demonstrates overall differences in root traits across biomes associated with the collaboration axis. Subtropical and tropical species generally have thicker roots with lower SRL than temperate species (Freschet et al. 2017, Ma et al. 2018), possibly reflecting the dominance of arbuscular-mycorrhizal (AM) canopy trees of tropical forests compared to ectomycorrhizal (EcM) dominance in temperate forests (Tedersoo 2017). Since AM fungi colonize the root cortex, their hosts tend to have thicker root tips with a larger cortex than EcM-associated species, where the fungal mantle develops outside the root (Brundrett 2002, Kong et al. 2014). Biomes may also exhibit root trait differences associated with the conservation axis. Temperate ecosystems with lower temperatures, slower nutrient cycles, and shorter growing seasons, may in turn filter for species with ‘slow’, robust root traits (*e.g*., high RTD) to conserve resources over the long term (Grime 1977, Reich et al. 1998, Aerts and Chapin III 2000, Reich 2014). Tropical forests, in turn, may provide overall more favorable abiotic environments with longer growing seasons, higher temperatures, and faster microbial-driven nutrient cycling that would select for ‘fast’ roots (high root N) with a strong competitive advantage for resource uptake. Alternatively, tropical plant species may be expected to have ‘slower’ roots than temperate species because of higher pressure from natural enemies, *e.g.*, soilborne pathogens (Delgado-Baquerizo et al. 2020), selecting for chemically-protected and less palatable roots with high RTD (Freschet et al. 2017, Xia et al. 2021) and lower root N concentrations (Freschet et al. 2017, Laughlin et al. 2021). Owing to these different biotic and abiotic selective forces, subtropical and tropical tree species may take up distinct positions in the global root economics space, which currently mostly describes variation in root traits in temperate, continental ecosystems.

Subtropical and tropical systems are often treated as single homogenous biomes. Overlooking the large variation in evolutionary trajectories and geographical and environmental characteristics they comprise (Townsend et al. 2008, Fujii et al. 2018) constrains our knowledge of root trait variation within the (sub)tropics and its implications for global diversity. As a first example, evolutionary differences may be large as the tropics comprise multiple phytogeographic regions, *i.e.,* phytogeographical delineation based on evolutionary relationships.

This variability leads to greater evolutionary distinctness within the tropics and co-occurrence of deeply diverged lineages within and across continents (Carta et al. 2022). Secondly, in the American tropics, the arrival of boreo-tropical plant species (*e.g.*, *Quercus* and *Salix*), large topographic variation resulting from the rise of the Andes (Antonelli 2021), and rapid plant radiations at high elevations (Madriñán et al. 2013) may further enlarge root trait variation within the region. Thirdly, several taxonomic groups evolved unique, contrasting belowground symbiotic traits within and across continents, reflected by the local dominance of canopy trees like Dipterocarpaceae in Southeast Asia or Fagaceae and Juglandaceae in Central and South America highlands breaking the perceived dominance of AM-associated species and N-fixers in the tropics (Steidinger et al. 2019). Fourthly, geological changes in Africa have resulted in ecosystem turnover from the typical moist forests into savannas and dry woodlands (Raven et al. 2020), with savannas currently covering 50% of the African continent (Osborne et al. 2018).

These examples illustrate how - across the tropics - different continents would have their own root trait expressions because of their unique biogeographic and evolutionary history (Comas et al. 2012, Echeverría-Londoño et al. 2018, Weemstra et al. 2023) to fill their belowground niche in these heterogeneous environmental gradients.

Environmental gradients *e.g.*, in nutrient and water availability, may also differ within the (sub)tropical biomes, shaping belowground plant strategies (Fujii et al. 2018, Cusack et al. 2021, Yan et al. 2022b). In Australia, the presence of non-mycorrhizal proteoid roots on severely P- impoverished soils is characteristic (Lambers et al. 2006), while adaptations to fires play another important role in root allocation and trait expression. Plants in Africa had higher median RTD and SRL than sites in Asia and the Neotropics, which was partly attributed to root morphological plasticity in response to frequent exposure to water stress in African systems (Addo-Danso et al. 2020), and within African tree species, seedlings from dry and moist ecosystems had distinct root trait syndromes (Boonman et al. 2020). In other words, across the “global (sub)tropics’’, as a response to contrasting environmental gradients, it is expected that variation in root trait expressions depends on species’ niches, such as the “typical” climatic or soil conditions of the species, *i.e.*, niche position, as well as the size of the species’ environmental space, *i.e.*, niche breadth (Vleminckx et al. 2023).

Here, we aim to expand our understanding of plant biodiversity by assessing the contributions of subtropical and tropical biomes – and the variation within – to the global functional diversity currently known belowground. Specifically:

1. We assessed the degree of interspecific covariation in traits related to the belowground collaboration and conservation axes among the three biomes. From temperate to tropical ecosystems, we hypothesized that (H1) along the collaboration axis, roots shift from a “do-it-yourself” to an “outsourcing” strategy because AM associations dominate in tropical biomes. Along the conservation axis, towards the tropics (H1a), roots shift from “slow” to “fast”, as overall environmental conditions become more favorable, or (H1b) roots shift from a “fast” to “slow” strategy towards the tropics due to elevated soil-borne pathogen loads.
2. We compared the degree of root functional diversity of temperate, subtropical, and tropical biomes, and hypothesized that (H2) belowground traits in subtropical and tropical biomes are more functionally diverse compared to the temperate biome.
3. We determined the contribution of subtropical and tropical biomes to the global root trait space currently dominated by temperate root data. Because different (a)biotic constraints prevail and select for distinct root trait expressions, we hypothesized that (H3) species from subtropical and tropical biomes would occupy distinct spaces in, and their inclusion would thereby expand the global root trait space.
4. Finally, we explored drivers of interspecific root trait variation within subtropical and tropical biomes. We hypothesized (H4a) that interspecific root trait variation in subtropical and tropical biomes is associated with different aspects of the species’ niche, *i.e.*, niche and niche breadth, suggesting that specific belowground strategies might offer an overall fitness advantage across climatic and soil conditions in these biomes. Further, we hypothesized (H4b) that interspecific root trait variation is shaped by biogeographic drivers, particularly evolutionary variation among continents.

## Materials and methods

### Species-level root trait data

We focused on the four key root traits that make up the conservation and collaboration axes in the root economics space (Bergmann et al. 2020): mean root diameter (mm), SRL (m g^-1^), RTD (g cm^-3^), and root N concentration (mg g^-1^). Because trait expressions and covariations generally differ between woody and non-woody species (Roumet et al. 2016, Weemstra et al. 2016), we further included information on plant woodiness. Root trait data and plant woodiness were extracted from the Global Root Trait (<u>GRooT</u>) database ver.2 (Guerrero-Ramírez et al. 2021), which combines root trait observations from the Fine Root Ecology Database ver.3 (FRED; Iversen et al. 2021), the Plant Trait Database ver.5 (TRY; Kattge et al. 2011, 2020), data mobilized by the Tropical Root Trait Initiative (<u>TropiRoot</u>), additional datasets and unpublished data from Ghana (unpublished, S.D.Addo-Danso). Because the root economics space focus on traits associated with belowground resource uptake, we excluded trait data from coarse and transportive roots (identified by the data authors and data contributors as roots with diameter > 2 mm) that play no or only a marginal role in acquiring soil resources (McCormack et al. 2015).

Individual observations with an absolute error risk value (*i.e.*, the number of mean standard deviations (across all species within a trait) from the respective species means provided by GRooT (Guerrero-Ramírez et al. 2021) higher than 4 were excluded before calculating species-level mean values. Data compilation resulted in a dataset consisting of 1035 species for which data on *all* four core traits were available. For the individual traits, our dataset comprised 2391, 2031, 1747, and 1886 species for which we had data on SRL, mean root diameter, root N concentration, and RTD, respectively.

### Climates and floras across continents

Data from GRooT were joined with the new version of the World Checklist of Vascular Plants (WCVP; Govaerts et al. 2021). Due to limited data from the tropics, we categorized the tropical biome by aggregating the “montane tropical”, “seasonally dry tropical”, and “wet tropical” climate categories based on the WCVP climate description. WCVP species-level data for continents were aggregated, *i.e.*, if a subtropical or tropical species occurred in “Northern America” and “Southern America”, it was assigned to “America”, and if the species occurred in “Asia Tropical” and “Asia Temperate”, it was assigned to as “Asia”. When subtropical or tropical species were present in more than one continent, *e.g.*, *Dodonaea viscosa* Jacq, occurring in “Africa”, “Asia”, “Australasia”, “America”, and “Pacific”, we classified it as “Pansubtropical or pantropical”, respectively. Because global root trait assessments are mostly based on temperate species (Freschet et al. 2017, Bergmann et al. 2020, Carmona et al. 2021), we used the temperate biome as a root trait space reference.

### Species occurrences and climatic and soil conditions

To ensure that all subtropical and tropical species with available data on the four root traits were included in our analyses, and to calculate their climatic and soil niches (see below), we obtained species’ occurrence data (coordinates) from the Global Information Biodiversity Facility (GIBF; accessed October 2022) using the ‘occ_data’ function in the rgbif package (Chamberlain and Boettiger 2017). Data on (sub)tropical species’ occurrence were restricted between latitudes 35°N and 35°S. Data from GIBF were cleaned by (i) removing fossil specimens and managed, introduced, invasive, and naturalized species records; (ii) omitting non-terrestrial coordinates; and (iii) excluding coordinates based on the vicinity of country, country capitals, province centroids, and biodiversity institutions (with a buffer of 2 km) to minimize the chance that coordinates of e.g., planted species outside their natural distribution range would be included using the CoordinateCleaner R package (Zizka et al. 2019).

Climatic data were obtained from Chelsa version 1.2 (1 km resolution) and included mean annual temperature (°C), mean annual precipitation (mm), precipitation in wettest and driest months, temperature seasonality (*i.e.*, the standard deviation of the monthly precipitation estimates expressed as a percentage of the mean of those estimates and the standard deviation of the monthly mean temperatures), and maximum and minimum temperatures in warmest and coldest months, respectively (Brun et al. 2022). Soil data were retrieved from the Harmonized World Soil Database (1 km resolution, 0-30 cm, Appendix S1: Figure S2), (Fischer et al. 2008) including variables that generally serve as indicators of soil properties resulting from abiotic and biological processes acting at longer timescales (Garland et al. 2021) (Appendix S1: Table S1). In addition, total soil phosphorus (P) data at a 0.5-degree resolution were retrieved from the Global Gridded Soil Phosphorus Distribution Maps (Yang et al. 2014), but as these data were unavailable for 20 (sub)tropical species and for 12% of the species occurrence observations (*i.e.,* 44,688) and were correlated with cation exchange capacity (Pearson r = 0.66, p-value < 0.001; Appendix S1: Figure S2). Thus, we did not include soil P data in our analyses.

### Phylogenetic tree

To investigate the influence of evolutionary history on trait variation in subtropical and tropical biomes, we built a phylogenetic tree using a dated mega-tree for vascular plants as the backbone (‘GBOTB.extended.tre’; Jin and Qian 2019), which merged trees from Smith and Brown (2018) and Zanne *et al.,* (2014). We resolved missing species with the most commonly used ‘Scenario 3’ in the ‘V.Phylo.Maker R’ package (Jin and Qian 2019). The phylogenetic tree was visualized using the ‘ggtree’ package (Yu et al. 2017, 2018, Yu 2020).

## Data analysis

### Interspecific (co)variation in root traits among biomes & root functional diversity

The multivariate covariation in root traits among species, *i.e.*, the root trait space, within and across temperate, subtropical, and tropical biomes, was visualized using a principal component analysis (PCA) based on logarithmically transformed and scaled species-level mean values of all four core root traits. Differences in the multivariate root trait space among biomes were tested using a Permutation Multivariate Analysis of Variance (Permanova) based on 999 permutations with ‘adonis2’ in the vegan package (Oksanen et al. 2019). In addition, for each of the four core root traits, mean and dispersion (*i.e.*, standard deviation, trait variation) values, as well as their confidence intervals (based on 0.025 and 0.975 quantiles), were calculated across species per biome using the ‘cwm’ and ‘cwd’ functions in the BAT package, assuming equal species abundances (Cardoso et al. 2015) and based on 999 repetitions controlling for species richness by randomly selecting 100 species from each biome. We considered biomes to differ in their root traits if confidence intervals (CI) were not overlapping.

We calculated functional diversity by building hypervolumes (*i.e.*, shape and volume of high-dimensional objects using a threshold kernel density estimate) of the first two axes of the root trait PCA (Appendix S1: Table S2) (Mammola and Cardoso 2020). Specifically, we randomly selected 100 species from each respective biome and built the hypervolume using the ‘hypervolume_resample’ function in the hypervolume package (Blonder et al. 2018). Based on each hypervolume (*i.e.*, 100 hypervolumes for each of the three biomes), we calculated functional richness (*i.e.*, the total volume of the functional space), evenness (*i.e.*, overlap between the calculated volume and a simulated volume with even distribution of traits, with 1 as perfect overlap), and dispersion (*i.e.*, the average distance between the centroid and a sample of stochastic points) using the BAT package (Cardoso et al. 2015).

### Contribution of subtropical and tropical biomes to the global root trait space

To assess the proportion of subtropical and tropical functional spaces relative to the global root economics space, we determined the proportion of the root trait hypervolume unique to the respective biome relative to the root trait hypervolume of all other biomes combined (excluding that of the biome of interest, *e.g.*, the unique contribution of tropical species compared to the functional space covered temperate and subtropical species together), using the ‘hypervolume_overlap_statistics’ function (Blonder et al. 2018). Because the number of species differed between each biome and the rest of the space (*i.e.,* excluding the species from the target biome), we standardized by comparing hypervolumes with the same number of species (*i.e.,* using the number of species present in the smaller hypervolume in the target biome or the rest of the space as a baseline). The mean value for uniqueness and its 95% confidence intervals were calculated based on 999 repetitions.

### Drivers of subtropical and tropical root trait variation

We determined if subtropical and tropical root trait variation were associated with species’ climate and soil niche position and niche breadth and/or variation in evolutionary histories among continents. We first built species-specific climatic and soil niches (position and breadth). For species niche *positions*, first, we carried out two PCAs using logarithmic or square-root transformed (when needed to meet assumptions of normality) and scaled environmental species mean values: one PCA for climatic and one PCA for soil variables in which each data point represents a single species (Appendix S1: Tables S3 and S4 and Figures S3 and S4). For the climatic PCA, the first axis (PC1_clim_) explained 59% of the total variation and was positively related to MAP, precipitation in the wettest and driest months, and minimum temperature in the coldest month, and is referred to as a precipitation axis (Appendix S1: Table S3 and Figure S2). The second axis (PC2_clim_) explained an additional 25% of the variation, with positive values strongly associated with higher MAT, and mean temperatures in the warmest month, *i.e.*, a temperature axis. For the soil PCA, PC1_soil_ explained 41% of the variation, with positive values predominantly associated with higher silt content and cation exchange capacity (CEC). The PC2_soil_ explained 34% of the variation, with positive values being mostly related to lower clay content (Appendix S1: Table S4 and Figure S4). Species’ climatic and soil niche *positions* were then extracted as the species’ scores on PC1 and PC2 of the climate and soil PCAs, respectively (Appendix S1: Figures S3 and S4). For species-specific niche *breadths*, we first carried out PCAs in which every point represents the environmental conditions associated with species’ occurrences, *i.e.*, multiple points for each species (Appendix S1: Figure S1), using logarithmic or square-root transformed (when needed to meet assumptions of normality) data. Subsequently, we built hypervolumes for each species based on the first two axes from the respective climatic and soil PCAs. For niche breadth, we used the ‘hypervolume_resample’ function in the ‘hypervolume’ package (Blonder et al. 2018).

Secondly, we calculated the phylogenetic relatedness of species between continents, *i.e.*, the evolutionary distance between the flora represented in our dataset from different continents. To do that, we followed the methodology described in Taylor *et al.,* (2023): first, we calculated standardized effects sizes of phylogenetic turnover (Phylo_SES_) between continents using the ‘phylobeta_ses’ function (Procheş et al. 2006) in the phyloregion package (Daru et al. 2020). We used Phylo_SES_ to correct for taxonomic turnover. Specifically, phylobeta_ses determines the observed phylogenetic turnover (Phylo_OBS_) between continents based on their corresponding phylogenies. It shuffles tip names in the phylogeny of each continent (1000 iterations) to create null assemblages to compute the mean phylogenetic turnover (Phylo_NULL_mean_) and its standard deviation (Phylo_NULL_sd_ ) (Taylor et al. 2023). Phylo_SES_ is then calculated as:

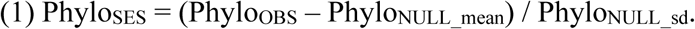

Next, we created a distance matrix using Phylo_SES_ values to carry out a Principal Coordinates Analysis (PCoA; Gower 1966) and extracted the first two PCoA axes representing the phylogenetic relatedness among continents, which accounts for most of the variation (Appendix S1: Figures S4 and S5). For the subtropical biome, differences in phylogenetic relatedness were overall associated with phylogenetic differences between Asia and Australasia, captured by the first principal coordinate axis (PCoA1_phyl_), and phylogenetic differences between pansubtropical (*i.e.,* species with a pansubtropical distribution were considered as from another “continent”) and Asia, Africa, America, and Australasia were captured by the second principal coordinate axis (PCoA2_phyl_, Appendix S1: Figure S5). For the tropical biome, differences in phylogenetic relatedness were overall associated with phylogenetic differences between Asia and America, captured by the PCoA1_phyl,_ and between pantropical and Asia, Australasia and America captured by the PCoA2_phyl_ (Appendix S1: Figure S6).

Finally, for each biome, we used linear models to determine if subtropical and tropical root trait variation (*i.e*, SRL, RTD, mean root diameter, and root N concentration) are associated with species’ climate and soil niche position (PC1_clim_ + PC2_clim_ + PC1_soil_ + PC2_soil_), niche breadth (Breadth_clim_ + Breadth_soil_), and species’ evolutionary histories (PCoA1_phyl_ + PCoA2_phyl_). To distinguish the effects of evolutionary history from other biogeographic aspects, we compared models including either phylogeny (PCoA1_phyl_ and PCoA2_phyl_) or continent using the Akaike Information Criterion (AIC). Evidence of the potential role of evolutionary history on root trait variation was associated with either the best model including phylogeny or when models including either phylogeny or continents had ΔAIC < 2. All data were analyzed in R Statistical Software ver. 4.1.1 (R Core Team 2023).

## Results

Species-level root trait data came from 1618 temperate, 341 subtropical, and 775 tropical species, representing 1418 woody, 1167 non-woody, and 41 woody/non-woody species, *i.e.*, plants whose woodiness is mixed or convoluted (Iversen et al. 2021). Of the woody species, 528 (37% of the total number of woody species) occurred in the temperate biome, 243 (17%) species in the subtropics, and 647 (46%) in tropical biomes. We first found that among the non-woody species, the vast majority (1007 species; 86% of all non-woody species) were temperate species, and only 73 (6%) and 87 (7%) species occurred in the subtropical and tropical biomes, respectively (Appendix S1: Figure 8). Owing to these data limitations on non-woody subtropical and tropical species (which largely represent the global coverage) and the potential of confounding effects when evaluating belowground functional biogeographic patterns, we restricted our results exclusively to the woody species, hereafter: ‘species’.

### Interspecific (co)variation in root traits among biomes and root functional diversity

The covariation in root traits among species, *i.e.*, multivariate root trait space, was captured by two main axes. The first component (PC1), recognized as the collaboration axis, explained 45% of the total interspecific variation in root traits, with SRL negatively and mean root diameter positively loading on PC1 (Figure 1). The second component (PC2), recognized as the conservation axis, explained an additional 36 % of the overall variation, with RTD negatively and root N concentration positively loading on PC2 (Figure 1). There was evidence for a shift in the collaboration axis (SRL and diameter) from do-it-yourself (high SRL) to an outsourcing strategy (high RD) from temperate to tropical biomes (Permanova: F= 33.3, p<0.001; Figure 1).

**Figure 1.**
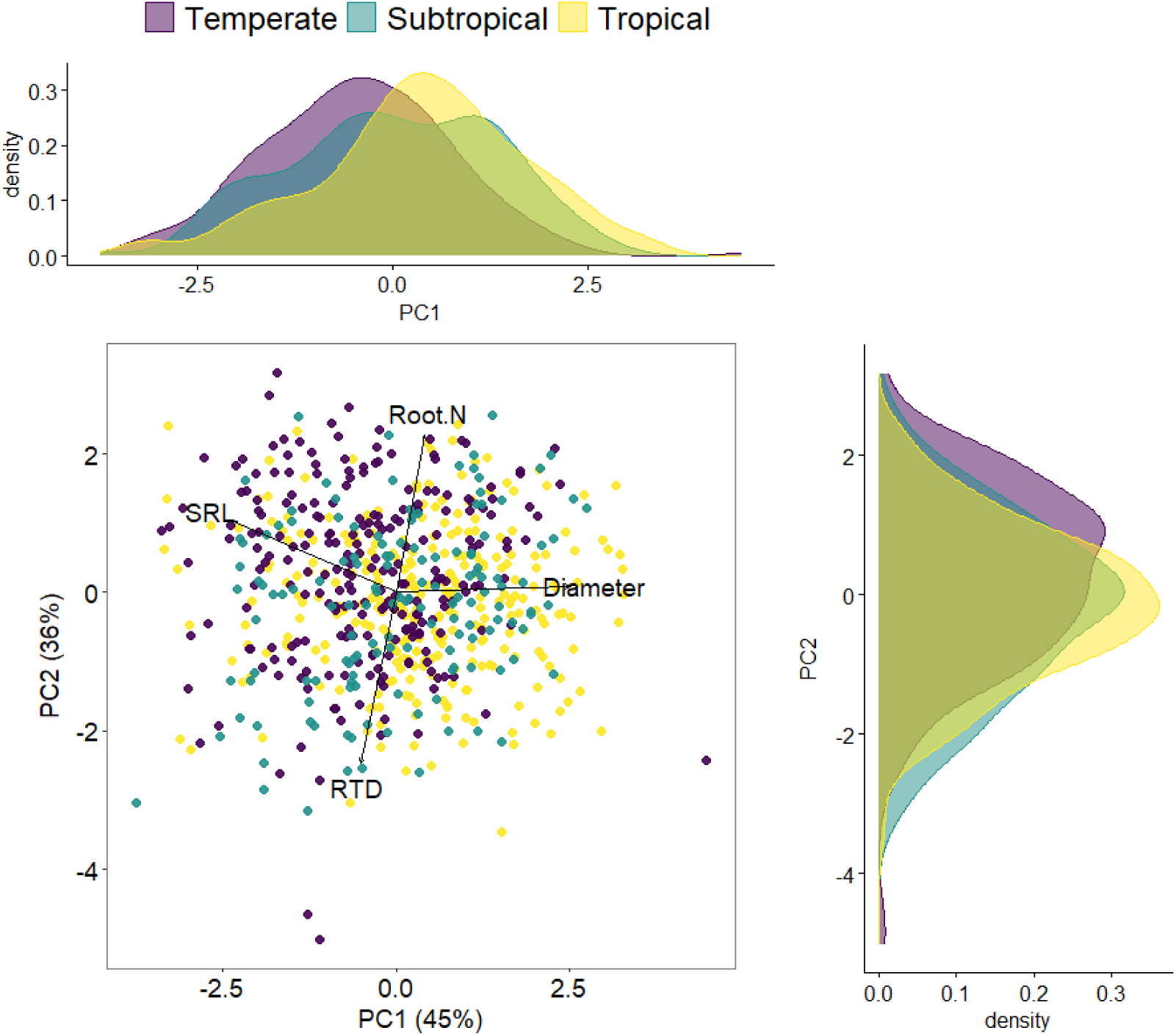
Variation in the root functional space for species among biomes. The root space is visualized using a principal component analysis (PCA). Traits included are mean root diameter, specific root length (SRL), root tissue density (RTD), and root nitrogen concentration (Root N). Each point represents a species, different colors refer to different biomes. Density distributions of the first two axes are shown on the upper and right sides. PCA and density distributions are based on 235, 143, and 289 species for temperate, subtropical, and tropical biomes, respectively. Bivariate correlations between traits across and within biomes are shown in Appendix S1: Figure S8.

Examining root trait variation for traits separately, we found evidence that temperate species had higher SRL and lower mean root diameter than subtropical and tropical species, but subtropical and tropical species were indistinguishable from each other (Figure 2, Table 1). Root N concentration was higher for tropical than subtropical species, with intermediate values for temperate species. Temperate species had lower RTD than subtropical and tropical species, while subtropical and tropical species had similar RTD. Overall, dispersion in the four individual traits did not vary significantly among biomes, except for RTD, which was less dispersed in tropical species than in temperate and subtropical species (Table 1). Further, there were no significant differences in functional richness, evenness, and dispersion among biomes when considering the conservation and collaboration axes together and controlling by species richness (Table 2).

**Figure 2.**
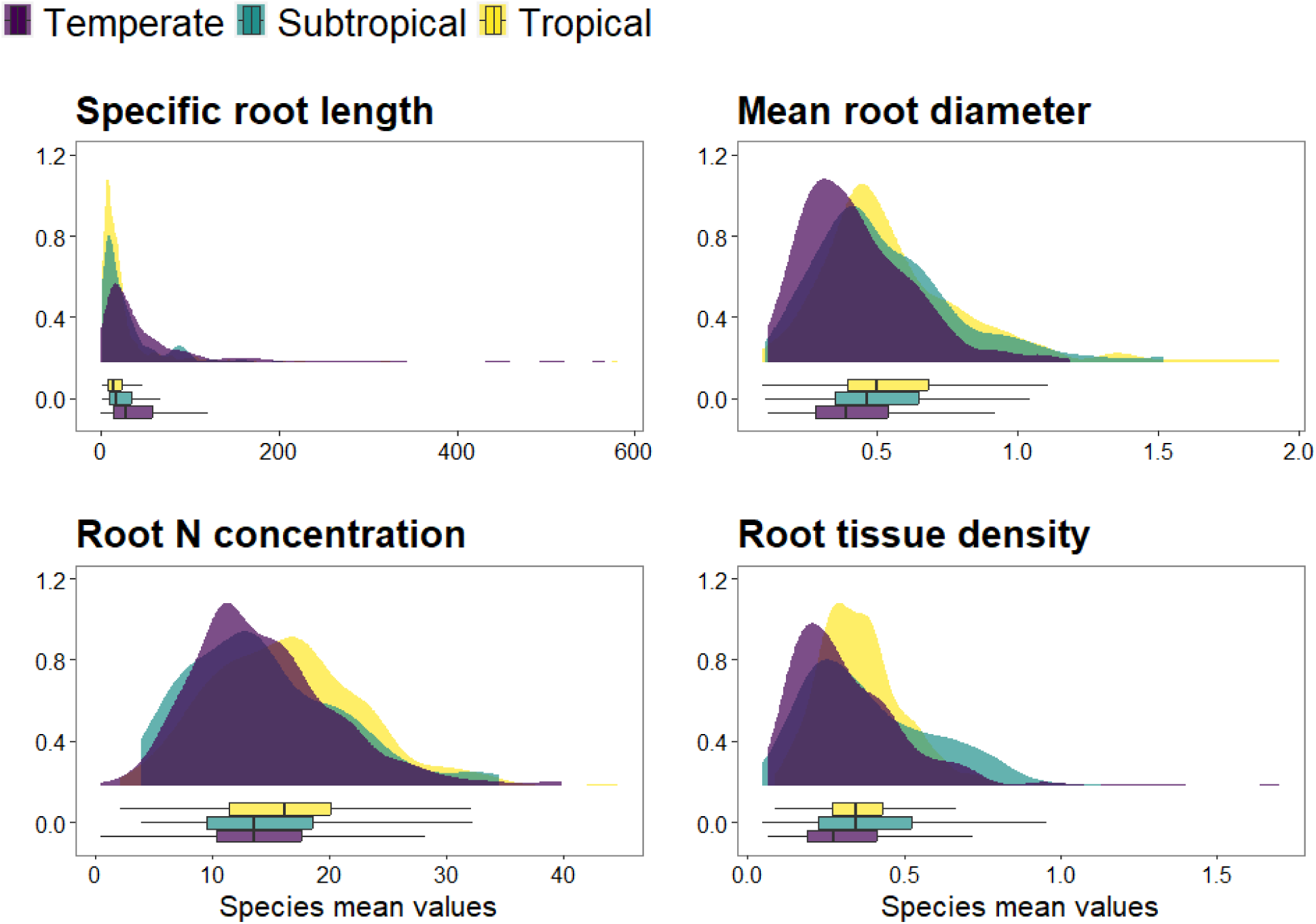
Histograms for species mean values for each of the target root traits. Traits included are specific root length (m g^-1^; n = 469, 233, and 615 temperate, subtropical, and tropical species, respectively), mean root diameter (mm; n = 353, 228, and 614 temperate, subtropical, and tropical species, respectively), root nitrogen (N) concentration (mg g^-1^; n = 416, 156 and 322 temperate, subtropical and tropical species, respectively), and root tissue density (g cm^-3^; n= 329, 223, and 598 temperate, subtropical, tropical species, respectively). Horizontal bar graphs show the medians (black, vertical lines) and boxes extending from the first and third quartile and the lines representing the min and max values that are not outliers of the trait distributions per biome.

**Table 1.** Mean and dispersion values for the four target traits. Based on logarithmically transformed species-level mean values for specific root length, mean root diameter, root nitrogen concentration, and root tissue density. Mean and dispersion values were calculated, standardizing species richness across biomes by randomly selecting 100 species for each biome. Mean and 95 % confidence intervals (CI, in gray) were based on 999 iterations. Different letters indicate differences among biomes based on overlapping confidence intervals (0.025 and 0.975%). Values were back-transformed with exponential.

| Traits |  | Temperate | Subtropical | Tropical |
| --- | --- | --- | --- | --- |
|  |  | -----Mean (95% CI) ----- |  |  |
| **Specific root length**<br>(m g<sup>-1</sup>) | Mean | 27.9<br>(22.8, 33.4)<sup>a</sup> | 17.1<br>(14.6, 19.8)<sup>b</sup> | 13.8<br>(11.6, 16.4)<sup>b</sup> |
|  | Dispersion | 3.0<br>(2.5, 4.0)<sup>a</sup> | 2.8<br>(2.5, 3.0)<sup>a</sup> | 2.6<br>(2.2, 3.0)<sup>a</sup> |
| **Mean root diameter**<br>(mm) | Mean | 0.39<br>(0.36, 0.41)<sup>a</sup> | 0.47<br>(0.44, 0.51)<sup>b</sup> | 0.51<br>(0.47, 0.55)<sup>b</sup> |
|  | Dispersion | 1.6<br>(1.5, 1.7)<sup>a</sup> | 1.6<br>(1.5, 1.7)<sup>a</sup> | 1.6<br>(1.5, 1.7)<sup>a</sup> |
| **Root nitrogen concentration**<br>(mg g<sup>-1</sup>) | Mean | 13.2<br>(12.1, 14.2)<sup>a,b</sup> | 13.0<br>(12.3, 13.6)<sup>a</sup> | 15.0<br>(13.9, 16.1)<sup>b</sup> |
|  | Dispersion | 1.6<br>(1.4, 1.8)<sup>a</sup> | 1.6<br>(1.5, 1.7)<sup>a</sup> | 1.5<br>(1.4, 1.6)<sup>a</sup> |
| **Root tissue density**<br>(g cm<sup>-3</sup>) | Mean | 0.28<br>(0.25, 0.30)<sup>a</sup> | 0.33<br>(0.30, 0.36)<sup>b</sup> | 0.34<br>(0.32, 0.36)<sup>b</sup> |
|  | Dispersion | 1.7<br>(1.6, 1.8)<sup>a</sup> | 1.8<br>(1.7, 1.9)<sup>a</sup> | 1.5<br>(1.4, 1.5)<sup>b</sup> |

**Table 2.** Functional richness, evenness, and dispersion for temperate, subtropical, and tropical biomes. Mean and 95 % confidence intervals (CI, in gray) based on 999 iterations. Different letters indicate significant differences among biomes based on non-overlapping confidence intervals (0.025 and 0.975%).

| Functional | Temperate | Subtropical | Tropical |
| --- | --- | --- | --- |
|  | ----- Mean (95% CI) ----- |  |  |
| Richness | 29.5<br>(24.9, 35.8)<sup>a</sup> | 31.9<br>(27.3, 36.9)<sup>a</sup> | 30.5<br>(24.8, 35.8)<sup>a</sup> |
| Evenness | 0.41<br>(0.24, 0.54)<sup>a</sup> | 0.52<br>(0.47, 0.59)<sup>a</sup> | 0.44<br>(0.38, 0.51)<sup>a</sup> |
| Dispersion | 2.10<br>(1.92, 2.41)<sup>a</sup> | 2.17<br>(1.98, 2.36)<sup>a</sup> | 2.14<br>(1.90, 2.39)<sup>a</sup> |

### Contribution of subtropical and tropical biomes to the global root trait space

We determined the unique root trait functional space of subtropical and tropical species relative to the global root trait space (Figure 3). Together, subtropical and tropical biomes accounted for at least 40% of the unique functional space. In other words, of the multidimensional root trait space across all three biomes, ∼37% is shared among biomes, 23% (CI: 15 – 30%) is unique to temperate, 23% to tropical (CI: 17 – 30%), and 17% (CI: 10 – 23%) to subtropical biomes.

**Figure 3.**
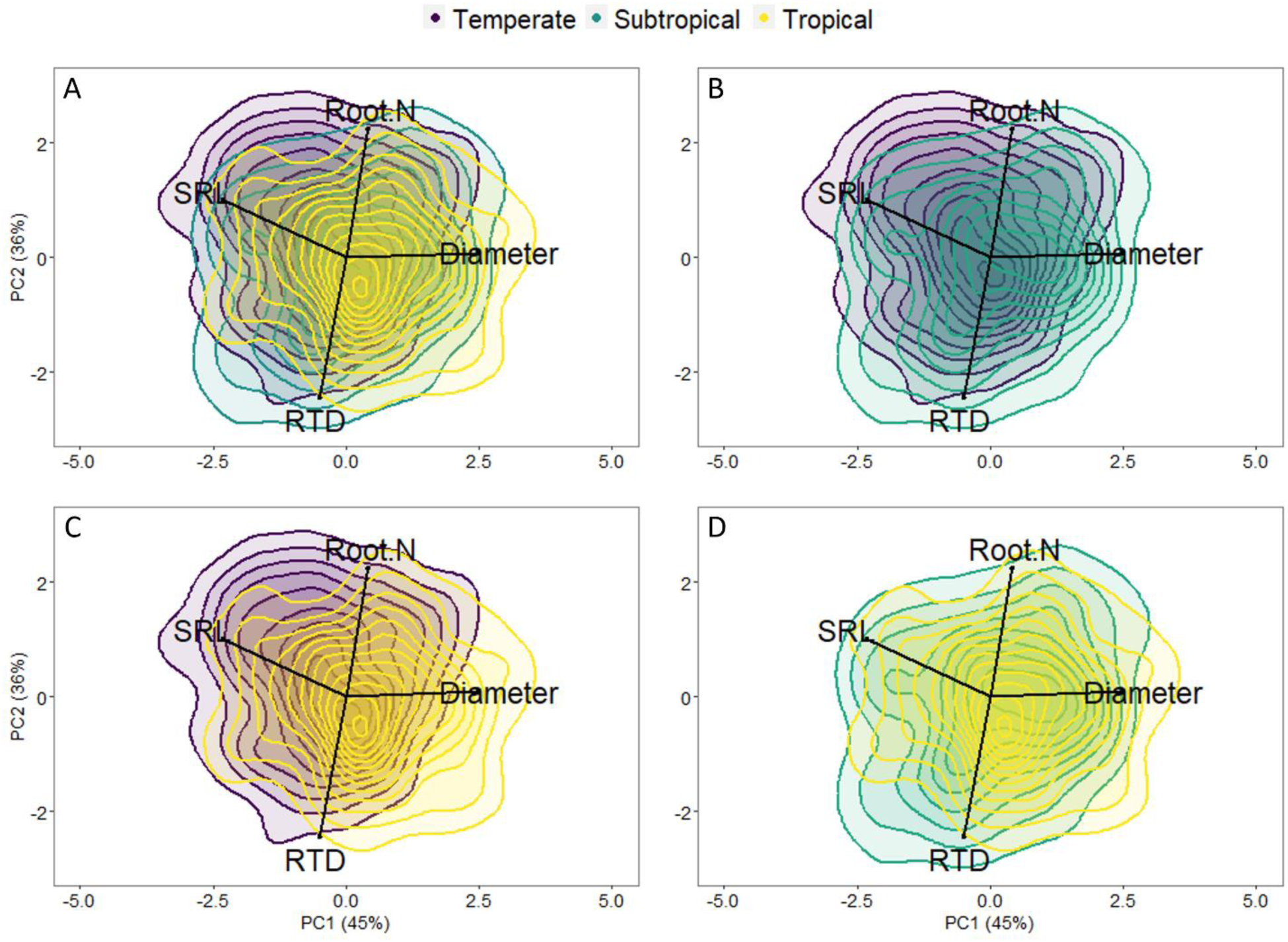
Principal component analysis acts as a graphical representation of the unique, i.e., unique multivariate strategies in each biome, and overlapping space, i.e., shared multivariate strategies between (A) all three biomes, (B), temperate and subtropical, (C) temperate and tropical, and (D) subtropical and tropical. The mean value for uniqueness and its 95% confidence intervals were calculated based on 999 repetitions. The traits included are root diameter, specific root length (SRL), root tissue density (RTD), and root nitrogen concentration (Root N).

### Drivers of subtropical and tropical root trait variation

The inclusion of evolutionary history either improved or performed similarly to subtropical models including continents, suggesting a role of evolutionary history in root trait variation (Appendix S1: Table S5). Subtropical models explained 27, 9, 4, and 56% variance for SRL, diameter, root N, and RTD respectively (Table 3). For instance, higher SRL values were associated with higher climatic niche breadth (Breadth_clim_) and species linked to environments with lower precipitation. Further, species from phylogenetic-related regions were similar in their SRL (*e.g.*, moving from higher values of SRL in Australasia species and lower species distributed across continents, *i.e.,* pansubtropical species, Figure 4). Higher RTD values were associated with species linked to environments with higher precipitation and were also similar among phylogenetic-related regions, with higher values for pansubtropical species and lower values in Australasia species (Figure 4). For the tropics, we found evidence of a potential role of evolutionary history for diameter and root N (Appendix S1: Figure S9), with differences for SRL and RTD associated with other biogeographic aspects captured by continents and not to evolutionary history per se (Appendix S1: Figure S10). Yet, the full models, including species niches and biogeographic/evolutionary drivers explained 10% or less variation (Table 3).

**Figure 4.**
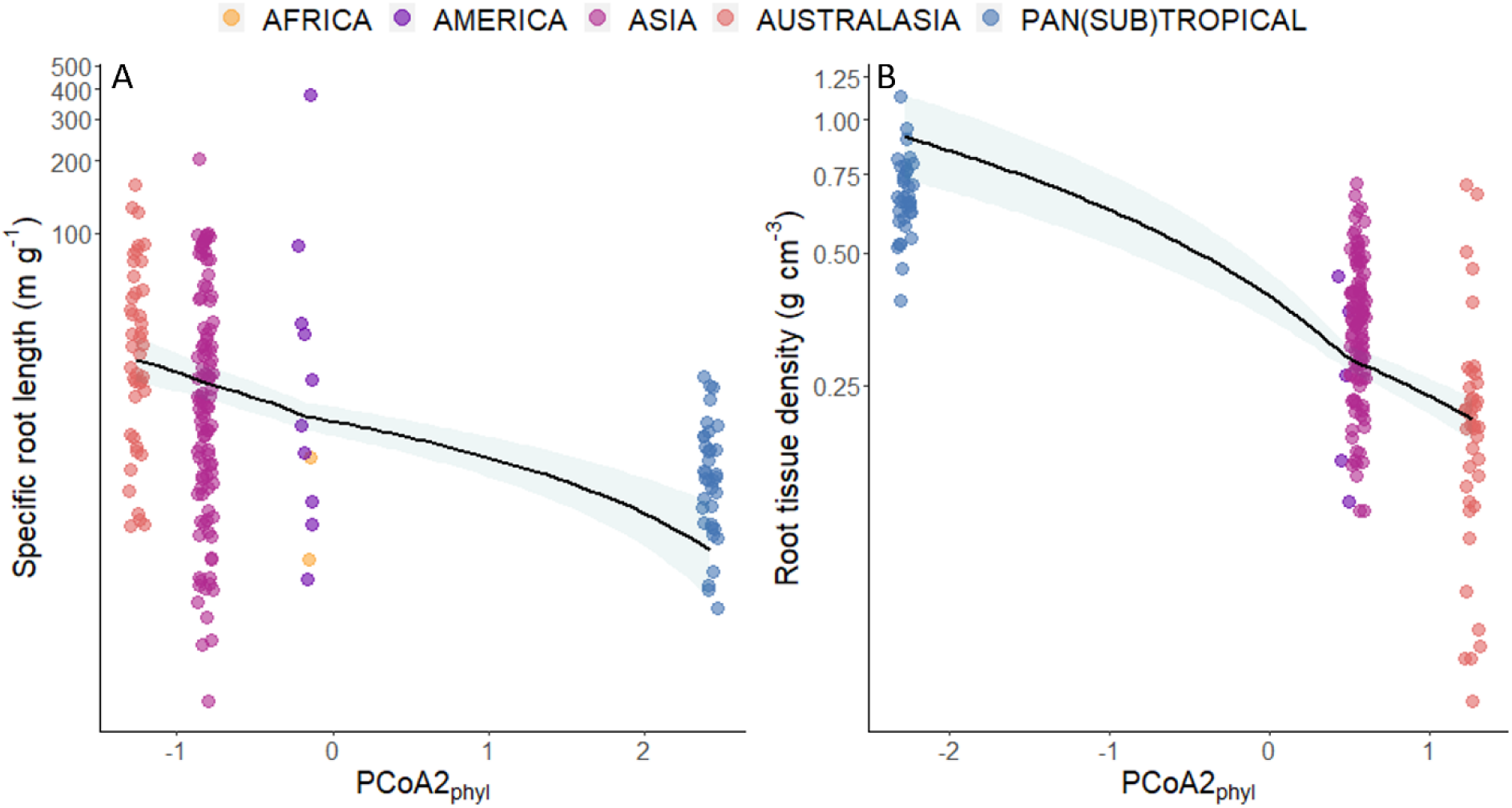
Phylogenetic relatedness (PCOA2_phyl_), i.e., evolutionary distance between the flora represented in our dataset from different continents, explaining interspecific variation in (A) specific root length and (B) root tissue density across subtropical species. Species from continents that are similar phylogenetically share similar specific root length and root tissue density values.

**Table 3.** Best models explaining root trait variation across species in subtropical and tropical biomes. To maximize the number of species included in the models, models for each trait varied in terms of species number and composition. *Df*, degrees of freedom; *F*, F statistic; and *P*, probability value. Bold *P* values represent significant relationships between root traits and species climate and soil niches (position and breadth) and biogeographic and evolutionary variables (ɑ = 0.05).

|  | Specific root length |  |  | Mean root diameter |  |  | Root N concentration |  |  | Root tissue density |  |  |
| --- | --- | --- | --- | --- | --- | --- | --- | --- | --- | --- | --- | --- |
|  | Df | F | P | Df | F | P | Df | F | P | Df | F | P |
| **Subtropical** |  |  |  |  |  |  |  |  |  |  |  |  |
| Breadth<sub>clim</sub> | 1 | ↑13.2 | **<0.001** | 1 | ↓5.68 | **0.018** | 1 | 0.13 | 0.723 | 1 | 0.47 | 0.493 |
| PC1<sub>clim</sub> | 1 | ↓13.7 | **<0.001** | 1 | 2.34 | 0.128 | 1 | 0.27 | 0.601 | 1 | ↑12.66 | **<0.001** |
| PC2<sub>clim</sub> | 1 | 0.06 | 0.806 | 1 | 0.47 | 0.492 | 1 | 2.16 | 0.144 | 1 | 1.08 | 0.300 |
| Breadth<sub>soil</sub> | 1 | ↓3.04 | 0.083 | 1 | 2.27 | 0.133 | 1 | 0.01 | 0.926 | 1 | 0.30 | 0.585 |
| PC1<sub>soil</sub> | 1 | 0.07 | 0.796 | 1 | 0.85 | 0.356 | 1 | 1.08 | 0.031 | 1 | ↑2.87 | 0.091 |
| PC2<sub>soil</sub> | 1 | 0.03 | 0.864 | 1 | 0.26 | 0.613 | 1 | 0.04 | 0.842 | 1 | 0.98 | 0.320 |
| PCoA1<sub>phyl</sub> | 1 | ↑3.90 | **0.049** | 1 | 0.13 | 0.722 | 1 | 0.25 | 0.619 | 1 | ↓4.32 | **0.039** |
| PCoA2<sub>phyl</sub> | 1 | ↓34.83 | **<0.001** | 1 | 0.18 | 0.671 | 1 | 0.53 | 0.467 | 1 | ↓88.85 | **<0.001** |
| Residuals | 186 |  |  | 179 |  |  | 124 |  |  | 174 |  |  |
| R<sup>2</sup> | 0.27 |  |  | 0.09 |  |  | 0.04 |  |  | 0.56 |  |  |
| **Tropical** |  |  |  |  |  |  |  |  |  |  |  |  |
| Breadth<sub>clim</sub> | 1 | ↑8.04 | **0.004** | 1 | ↓5.86 | 0.015 | 1 | ↑4.62 | **0.032** | 1 | 2.00 | 0.158 |
| PC1<sub>clim</sub> | 1 | ↓13.36 | **<0.001** | 1 | 1.58 | 0.209 | 1 | 2.13 | 0.145 | 1 | ↑5.26 | **0.022** |
| PC2<sub>clim</sub> | 1 | 0.51 | 0.477 | 1 | 0.002 | 0.962 | 1 | 1.06 | **0.030** | 1 | 0.003 | 0.960 |
| Breadth<sub>soil</sub> | 1 | 1.53 | 0.217 | 1 | 0.35 | 0.553 | 1 | 0.73 | 0.392 | 1 | 0.05 | 0.820 |
| PC1<sub>soil</sub> | 1 | 0.05 | 0.827 | 1 | ↑3.44 | 0.064 | 1 | 0.80 | 0.373 | 1 | ↓5.65 | 0.017 |
| PC2<sub>soil</sub> | 1 | ↓3.02 | 0.082 | 1 | ↑3.43 | 0.064 | 1 | ↓6.16 | **0.013** | 1 | 0.002 | 0.960 |
| Continent | 4 | 2.44 | **0.046** |  |  |  |  |  |  | 4 | 4.28 | **0.001** |
| PCoA1<sub>phyl</sub> |  |  |  | 1 | ↑3.47 | 0.063 | 1 | ↑13.47 | **<0.001** |  |  |  |
| PCoA2<sub>phyl</sub> |  |  |  | 1 | 0.13 | 0.715 | 1 | ↑8.80 | **0.003** |  |  |  |
| Residuals | 571 |  |  | 565 |  |  | 289 |  |  | 545 |  |  |
| R<sup>2</sup> | 0.09 |  |  | 0.05 |  |  | 0.09 |  |  | 0.09 |  |  |

## Discussion

Subtropical and tropical biomes are major contributors to global plant biodiversity (Cai et al. 2023). This extensive taxonomic diversity is associated with widespread functional diversity aboveground but also likely largely belowground, owing to the vast variability in belowground strategies of plants to acquire a multitude of resources from a heterogeneous and complex soil matrix (Weemstra et al. 2016). Our study shows how tropical woody species have different root traits compared to woody species in the temperate biome (*e.g.,* roots are overall thicker, less dense, and higher in N concentrations). Accounting for their unique combinations of root traits, subtropical and tropical species substantially expanded the global belowground functional space. Even though biomes have similar functional diversity and partially overlap in root functional space, (sub)tropical species seem to employ distinct belowground strategies that have so far not been observed or are rare in the temperate biome.

### Root trait expressions differ between (sub)tropical and temperate species

Tropical and subtropical species both had significantly lower SRL and larger root diameters than temperate species, supporting the expected shift along the collaboration axis (from do-it-yourself to outsourcing) between biomes (Freschet et al. 2017, Ma et al. 2018). Investing in thick roots that typically have large cortices (Kong et al. 2014) and that permit high AM colonization rates (Brundrett 2002, Ma et al. 2018) may be a particularly relevant foraging strategy in tropical soils (Valverde-Barrantes et al. 2021) where soil P is generally limiting (Reich and Oleksyn 2004, Vitousek 2004) and can be more efficiently acquired by AM fungi than soil N (Smith and Read 2008), generally limiting in temperate soils. The overall thicker roots in the (sub)tropics may further be explained by the greater abundance of more basal superorders in the tropics *e.g.*, Magnoliids, that rely heavily on AM fungi (Valverde-Barrantes et al. 2015, 2016) than recently derived superorders at higher latitudes, such as Rosids and Asterids, that have thinner roots and are less dependent on AM symbiosis (Comas et al. 2012, Ma et al. 2018). However, tropical forests often are still dominated by Rosid (Euphorbiaceae, Moraceae, Fabaceae, etc) and Asterid (Sapotaceae, Rubiaceae, Lecythidaceae) rather than Magnoliid (except Lauraceae) families.

Therefore, greater dominance of basal clades may explain only a small part of the shift towards the outsourcing strategy in (sub)tropical species. Further, in tropical forests, large root trait diversification still occurs *within* families, while traits (*e.g.,* root diameter) converge *across* families (Weemstra et al. 2023).

Tropical species had, overall, both higher root N concentration and higher RTD than temperate tree species, which contrasts previous work (e.g., Gu *et al.,* (2014), who found lower RTD in tropical than temperate tree species) and corroborates others reporting higher root N in the tropics, but including non-woody species (Freschet et al. 2017, Ma et al. 2018). These results call for careful interpretation of shifts in individual traits. For example, we observed a tradeoff between RTD and root N corresponding to the conservation gradient across all species combined and when considering biomes separately. However, a higher root N in tropical species does not necessarily imply a lower RTD, nor can it be interpreted as a “faster” belowground strategy relative to temperate species if RTD simultaneously increases too. Improving our understanding of plant belowground functioning requires the establishment of stronger linkages among traits (e.g., RTD and root N), between traits and functions (e.g., root N and resource uptake) and their mechanistic underpinnings.

From a resource economics perspective, an increase in RTD towards the tropics suggests that low root turnover rates would be advantageous in environments with tropical climates to maximize the return on C investment over the long-life span of the roots. A higher RTD in warmer climates was also observed, mainly in temperate forests, by Laughlin *et al.,* (2021), suggesting that this pattern is consistent within and across biomes and it may be associated with species’ tolerance to freezing conditions. Species that successfully migrated to higher latitudes where freezing events occurred are - among others - characterized by producing cheap aboveground tissue with annual turnover (Zanne et al. 2014), and a similar evolutionary trajectory may have occurred belowground where temperate (and boreal) species produce cheap, low-RTD roots to minimize resource losses as roots are shed during winter. Currently, the assumed resource economics tradeoff between RTD and root lifespan is not or weakly supported by (only few) empirical data across temperate tree species (Withington et al. 2006, McCormack et al. 2012), but it may be more pronounced in (sub)tropical biomes where the lack of freezing events may permit longer root lifespans and greater returns of investing in more expensive, dense root tissue. Biotic drivers may further explain the denser roots of tropical species, as high RTD offer structural and/or chemical protection to roots against herbivores and soilborne pathogen loads (Xia et al. 2021) that are generally higher in tropical soils than at higher latitudes (Delgado-Baquerizo et al. 2020).

### (Sub)tropical biomes add unique belowground strategies to the global space

Mapping the observed multivariate root trait combinations in a global functional space firstly revealed substantial overlap (37%) between species across biomes. These results suggest that at a broad scale, common belowground trait syndromes - made up of these four traits - occur regardless of climate and bioregion, owing to physiological (*e.g.,* resource economics) and evolutionary constraints that plants across bioregions are subjected to (Reich et al. 2003).

Despite these common pressures, however, adding the root traits from species in subtropical and tropical biomes extended the global root trait space, so that (sub)tropical species occupied at least 40% of the unique multivariate trait combinations within this space. Our analyses accounted for different numbers of species across biomes, so that the unique belowground traits space of (sub)tropical species cannot be attributed to larger species numbers. Furthermore, the degree of functional diversity was similar across biomes, suggesting that different but not more functionally rich trait combinations were observed among (sub)tropical species. Possibly, (sub)tropical species are subjected to largely different abiotic and biotic environmental conditions selecting for different root trait combinations unadaptive in other environments, that are further discussed in the following section. The unique contributions of (sub)tropical species to the global belowground functional space, nonetheless, illustrates that a large part of our understanding of belowground resource strategies has so far remained elusive, and that accounting for (sub)tropical species (of which our study still covers only a fraction) is pertinent to obtaining a more complete picture of how trees function belowground in a variety of ecosystems.

### Evolutionary history and species’ niches explain root trait variation in subtropical biome

Evolutionary history and species niches explained root trait variation among subtropical species for SRL and RTD, suggesting that biogeographic imprints on SRL and RTD in the subtropics are associated with evolutionary history. In other words, subtropical species from continents that are more similar phylogenetically, tend to have similar SRL and RTD values, suggesting both the role of evolutionary history and phylogenetic conservatism in root trait variation. In contrast, there was only a weak or no detectable role of evolutionary history for the other traits in the subtropics and for any of the four traits in the tropics. These findings support weak phylogenetic effects on root trait variation previously reported across 218 neotropical tree species that may be ascribed to large trait diversification and niche differentiation *within* tropical plant families (but see: Pierick et al. 2021, 2023, Weemstra et al. 2023). Together, our results point toward phylogenetic conservatism being context-dependent, varying across biogeographical, phylogenetic scales (Weemstra et al. 2023) and across traits describing both the collaboration and conservation axes. For example, we found evidence of the influence of evolutionary history for RTD but not for root N concentration in the subtropics (and similarly for SRL and not for diameter). Interestingly, lower values of SRL and higher values of RTD were associated with species with cosmopolitan biogeographic distributions (i.e., pansubtropical species), likely conferring the ability to explore the soil matrix via mycorrhiza and exhibit dense roots, with these strategies needed for complex mutualistic- and antagonistic-biosynthesis costs (Xia et al. 2021).

Determining whether the lower explanatory power of the models explaining root trait variation in the tropics is related to ecological and evolutionary reasons or data availability remains challenging. Regional studies have highlighted that the role of soil niches in shaping root strategies is nutrient-dependent, with weak significant relationships between SRL and species niches based on soil calcium but not for other soil N, P and potassium (K) in the Amazon (Vleminckx et al. 2023). In addition, biotic factors like mycorrhizal interactions and soilborne pathogen loads and herbivory may have a bigger effect on root selection in the (sub)tropics, while environmental conditions such as freezing temperatures seem more important selecting traits in the temperate biome. While our findings support the growing evidence suggesting a weak role of overall soil conditions (at least at larger scales) and phylogeny explaining root trait variation in the tropics, this needs to be interpreted with caution because of data shortfalls.

### Moving forward belowground functional biogeography

Based on our empirical cross-continental findings, we propose to broaden our view of (belowground) biogeography by including different ecosystems, traits, and species, specifically:

- *Underrepresented (sub)tropical ecosystems*: within the tropics, root data from certain, adverse ecosystems (like savannas, dry forests, and high montane ecosystems) are even more scarce than from e.g., moist tropical forests. Under their unfavorable environmental conditions, even more distinct belowground adaptation may be observed and the global belowground traits space further expanded.
- *Additional traits*: (sub)tropical species may diverge in traits outside the four core traits proposed (Bergmann et al. 2020), depending on their environment. For example, in tropical, P-limited soils, root exudation rates (Dallstream et al. 2022), phosphatase activity (Guilbeault-Mayers and Laliberté n.d.), branching (Yan et al. 2022a, Weemstra et al. 2023), and mycorrhizal traits may be more relevant than e.g., or SRL. Broadening our scope beyond the temperate zone might require measuring different traits functionally relevant in other biomes.
- *Non-woody species:* herbaceous species are well represented in tropical and subtropical biomes (24-34% herbaceous species; Taylor et al. 2023), but are largely absent in root trait databases and, subsequently, global analyses and resulting conceptual frameworks. Their poor coverage likely biases our knowledge of ecological and evolutionary drivers behind species-level root trait variation at global scales (Ma et al. 2018), with potentially misleading assessments on the role of families shaping root trait variation across biomes (Carmona et al. 2021). For example, some families, such as Euphorbiaceae or Fabaceae, are largely sampled as trees in the tropics but as herbs in temperate areas, making it difficult to distinguish the effects of plant growth habits from ecological adaptations to different biomes on belowground trait expressions and variability (Valverde-Barrantes et al. 2020). These biases need to be considered when interpreting other global analyses and addressed in sampling and data mobilization efforts.
- *Environmental information at a relevant spatial scale*: Owing to the large heterogeneity of soils at small spatial scales (Ettema and Wardle 2002), root traits are likely to respond to small-scale soil environmental variation (Weemstra and Valverde-Barrantes 2022, Pierick et al. n.d.). The use of global soil dataset with a low spatial resolution in global analyses, as ours, may therefore incur a mismatch between the scale at which soil variables vary and roots respond and lead to weak relationships between edaphic factors and root trait variation. Accurately linking the soil environment to root trait expressions and predicting how plants respond and adapt to soil environmental variation may therefore require more small-scale (in situ) data collection prior when, for instance, aiming to disentangle drivers behind larger scale patterns.

## Conclusions

Our findings illustrate the unique contribution of subtropical and tropical species to the global root economics space, expanding it with belowground strategies that are either rare or absent in the temperate biome and strengthening the foundations of functional belowground biogeography. This study significantly improves our understanding of global belowground strategies using a cross-continental, cross-biome representation of underrepresented species and ecosystems and by elucidating key shortfalls that need to be considered in global analyses. As such, our work highlights the importance of including (sub)tropical species in conceptual models of root functional diversity to develop a more complete view of the various belowground strategies that underlie plant functional biogeography and biodiversity globally.

## Author contributions

All authors conceived and developed the idea as part of the Belowground Functional Biogeography theme of the Tropical Root Trait Initiative. N G-R compiled the data and developed the analyses with the input of MW, MNU, and CF. N G-R and MW led the development and writing of the manuscript, with substantial input from all authors.

## Acknowledgments

This study emerged as a collaborative effort of scientists interested in addressing questions about tropical root ecology and evolution: the Tropical Root Trait Initiative (TropiRoot). We thank New Phytologist for supporting this Initiative and our work through its 28^th^ Workshop: Coordinating and Synthesizing Tropical Forest Root Trait Studies: Understanding Belowground NPP, Root Responses to Global Change, and Nutrient Acquisition Dynamics across tropical forests, held at the Smithsonian Tropical Research Institute in Panama. We thank Sarah Sophie Weil and Kevin Mganga as friendly reviewers and Amanda Taylor for the code to calculate the phylogenetic turnover. NR.G-R thanks the Dorothea Schlözer Postdoctoral Programme of the Georg-August-Universität Göttingen and the DFG, grant number 316045089/GRK 2300 for their support. DF.C participation was supported by US Department of Energy (DOE) Office of Science Early Career Award DE – SC0015898, and US National Science Foundation (NSF) Division of Environmental Biology (DEB) Long Term Research in Environmental Biology (LTREB) award #2332006. LFL acknowledges the Bavarian State Chancellery (Project Amazon-FLUX).

## Supporting Information

**Appendix S1.** Supplementary Figures and Tables

## Supplementary Figures

**Figure S1.**
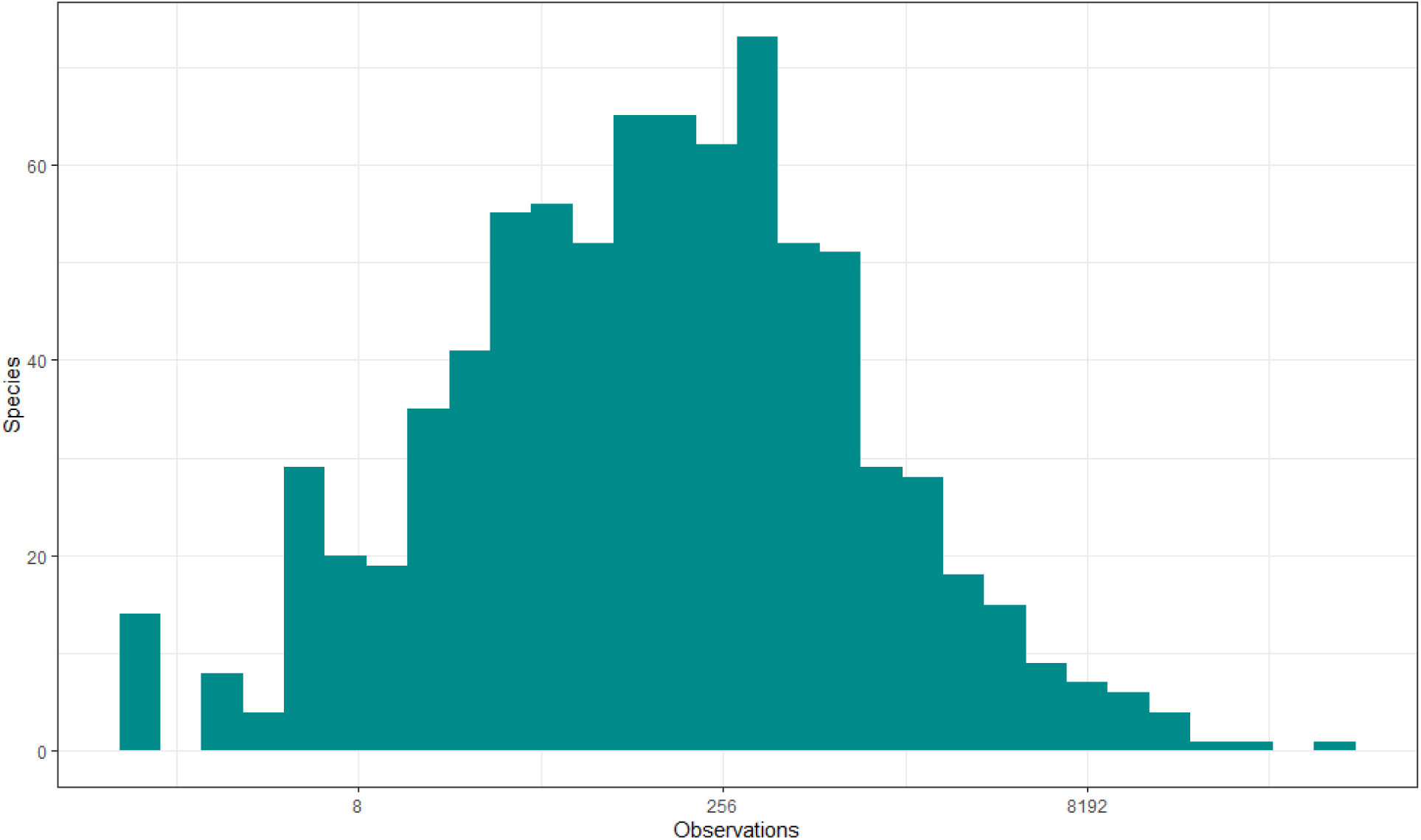
Number of coordinates (*i.e.*, observations) used to quantity species-specific niches.

**Figure S2.**
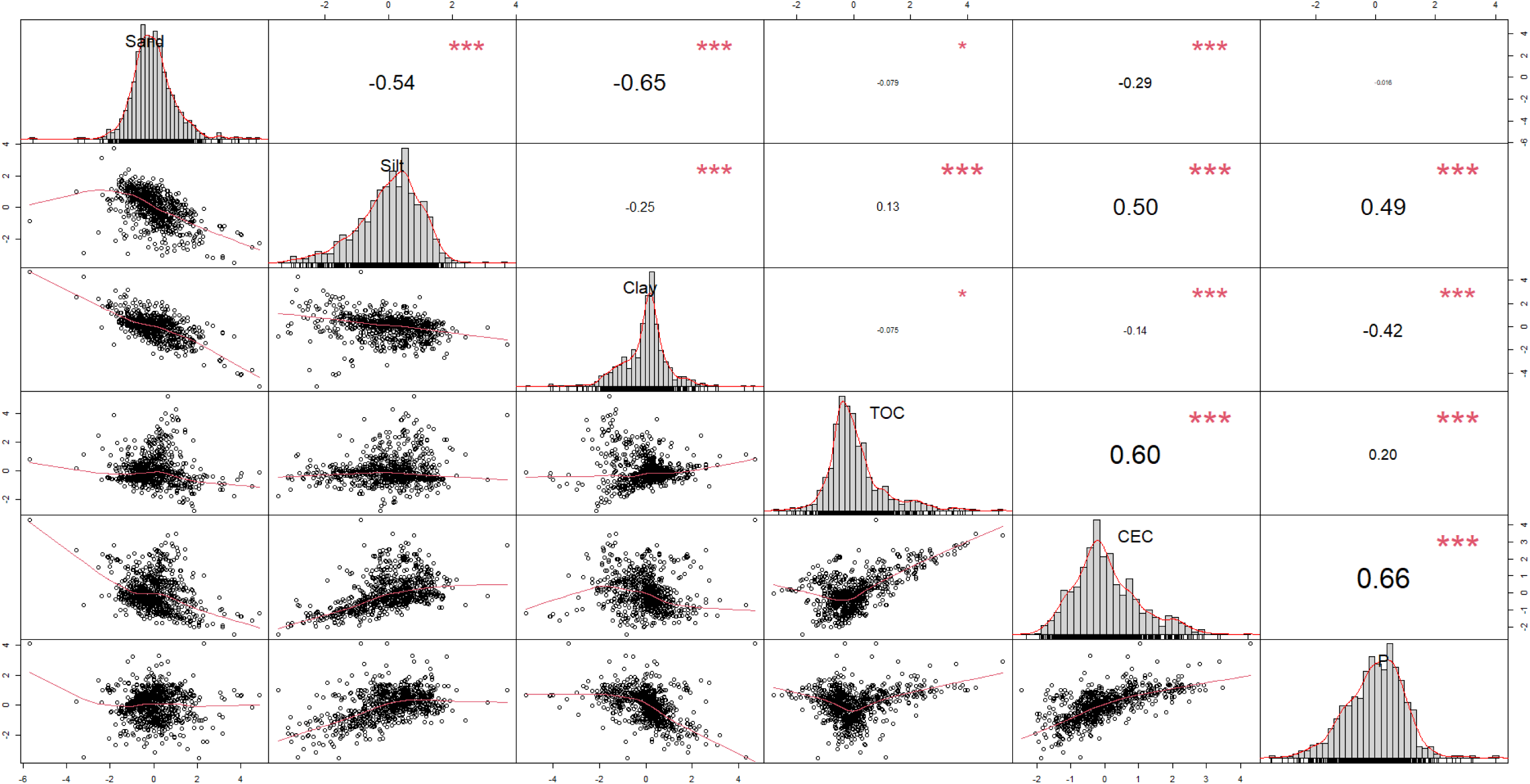
Pearson correlations for species-specific soil niches (position). Soil variables included sand, silt, and clay content, total organic carbon (TOC), cation exchange capacity (CEC), and total soil phosphorus (P).

**Figure S3.**
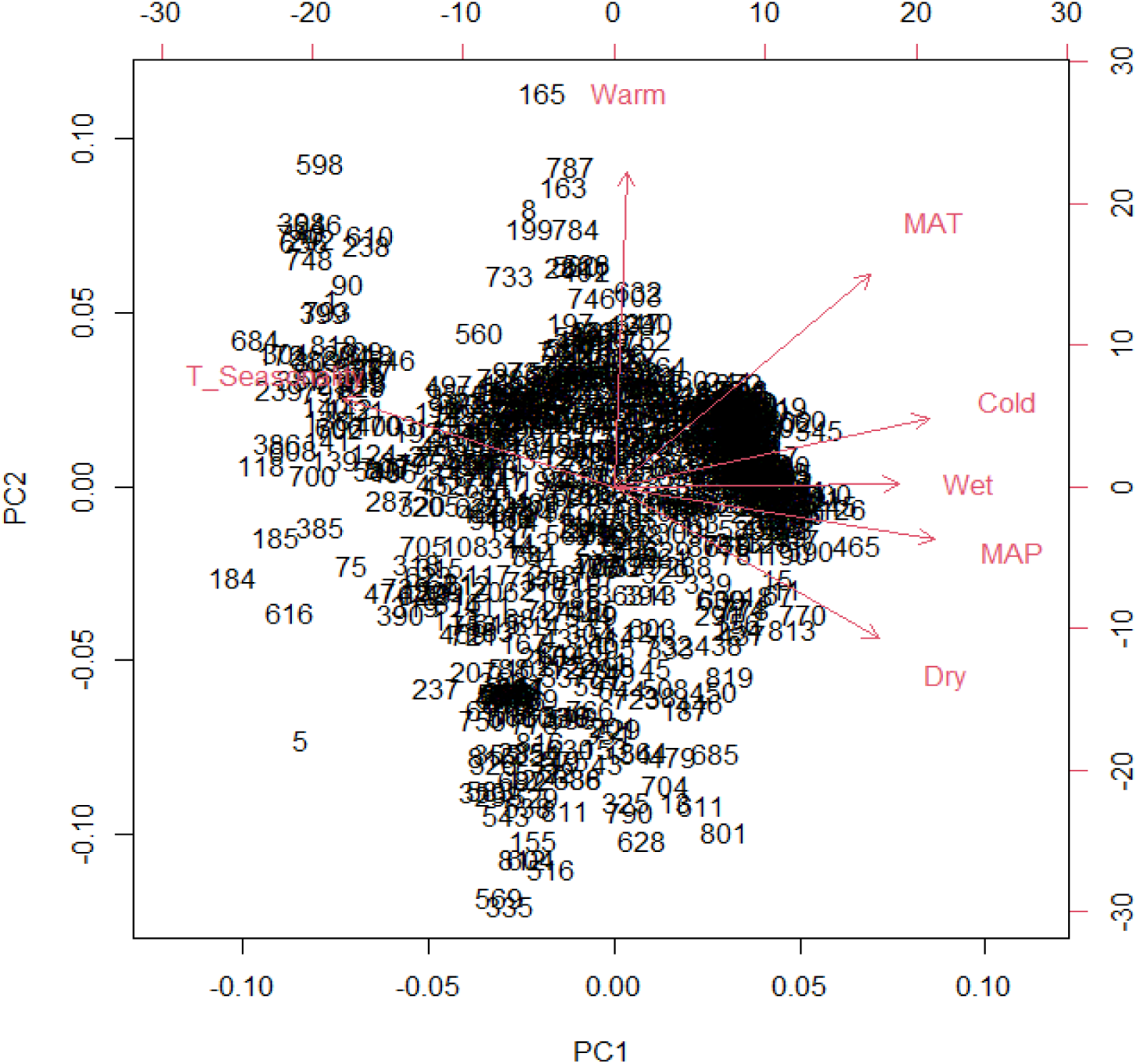
Principal component analysis of climatic variables for quantifying climate niche positions. Climatic variables include mean annual temperature (MAT, °C), mean annual precipitation (MAP, mm), precipitation in wettest and driest months (Wet and Dry), temperature seasonality (T_Seanonality), and maximum and minimum temperatures in warmest and coldest months (Warm and Cold), respectively.

**Figure S4.**
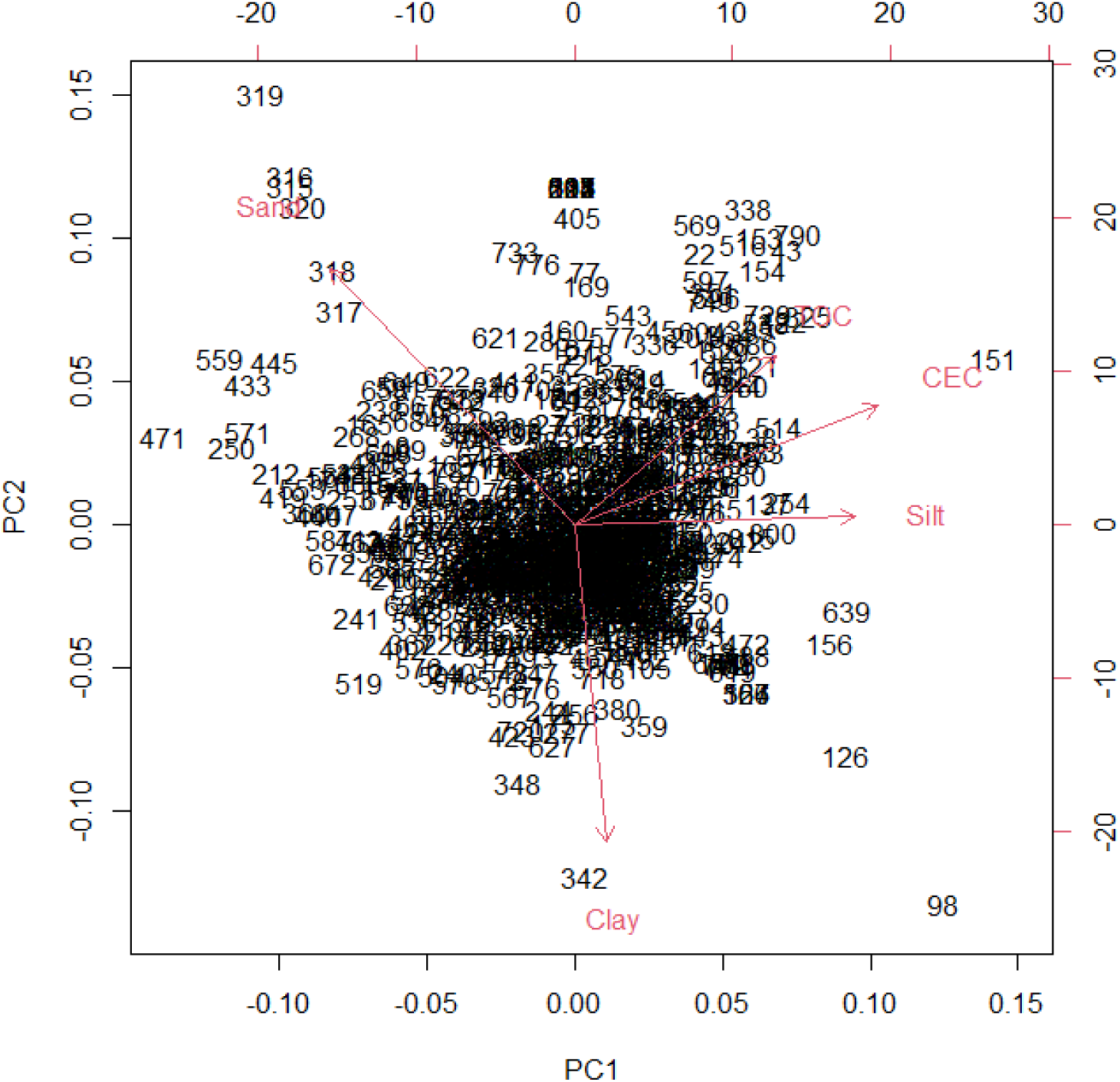
Principal component analysis of soil variables used to quantify soil niche positions. Soil variables includes topsoil sand, silt, and clay fraction (% wt), topsoil organic carbon (TOC, % weight), and topsoil cation exchange capacity (CEC, cmol/kg).

**Figure S5.**
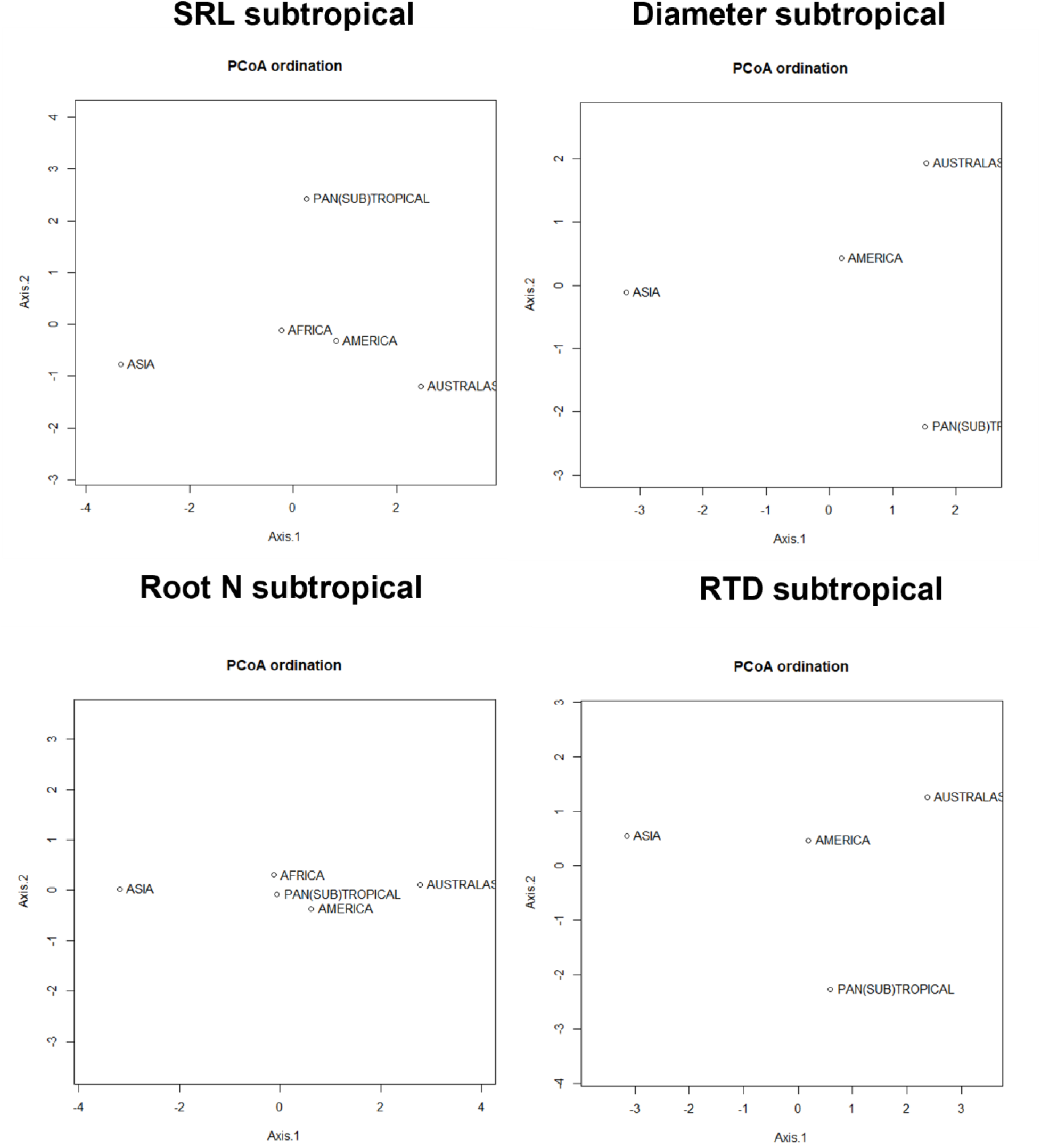
Principal Coordinate Analysis (PCoA) to determine the evolutionary distance between the subtropical flora represented in our dataset from different continents. The PCoA is based on distance matrices using standardized effects sizes of phylogenetic turnover.

**Figure S6.**
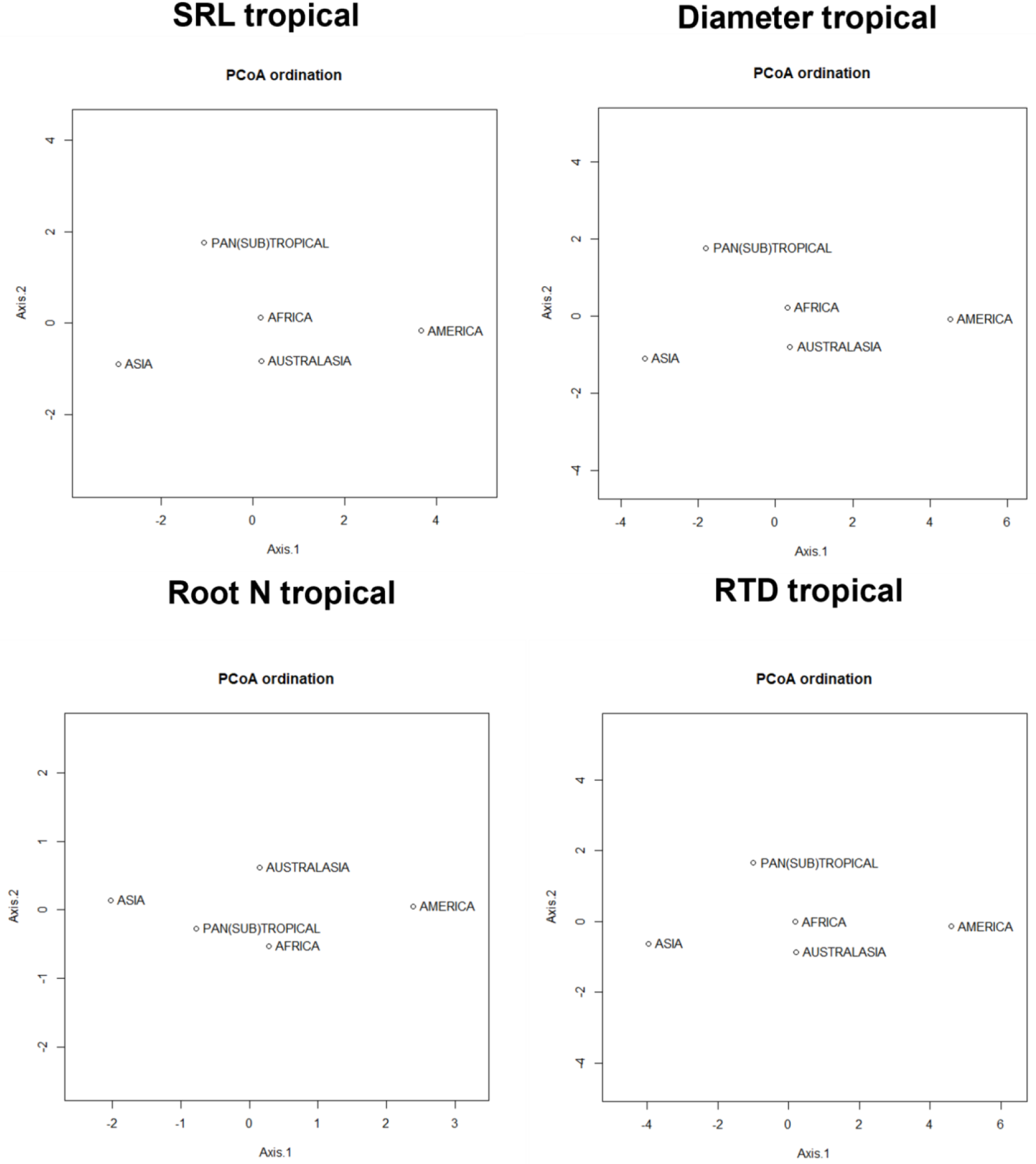
Principal Coordinate Analysis (PCoA) to determine the evolutionary distance between the tropical flora represented in our dataset from different continents. The PCoA is based on distance matrices using standardized effects sizes of phylogenetic turnover.

**Figure S7.**
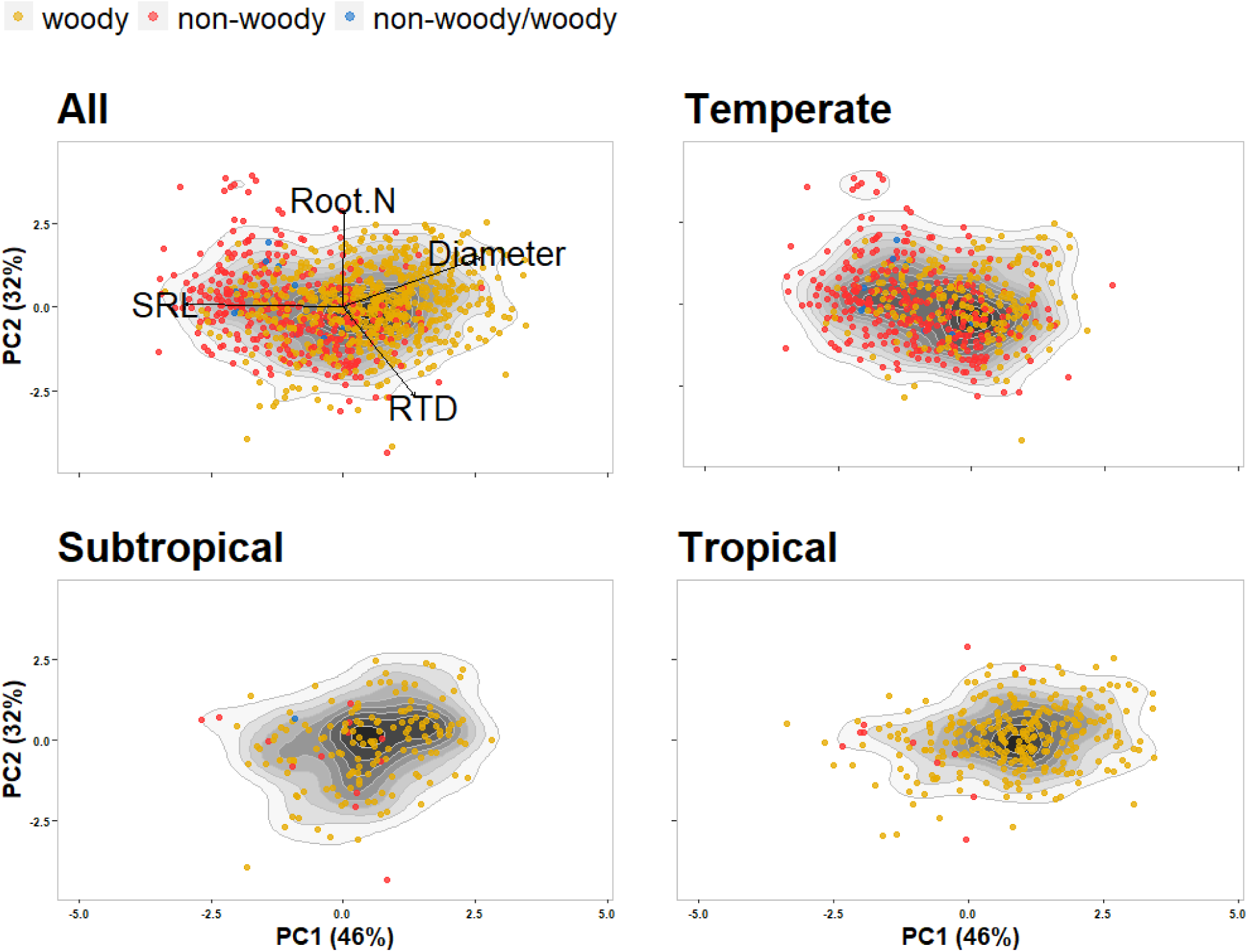
Root functional space including all available species mean values (n= 1035) together and separated into the three distinct regions (n= 561, 166, 308 for temperate, subtropical, and tropical species, respectively). Traits included are root diameter (mm), specific root length (SRL; m g^-1^), root tissue density (RTD; g cm^-3^), and root nitrogen concentration (Root N; mg g^-1^). Root functional spaces are visualized using a principal component analysis (PCA), with PC1 representing the collaboration gradient (SRL and Diameter) and PC2 the conservation gradient (Root N and RTD). The root functional space was separated by biomes to improve visualization but temperate, subtropical, and tropical panels represent the same PCA shown for all species together. Each point represents a species, with the colors representing woodiness (yellow dots = 667 woody, red dots = 320 non-woody, and blue dots = 10 non-woody/woody species). Contours are built using kernel density estimation.

**Figure S8.**
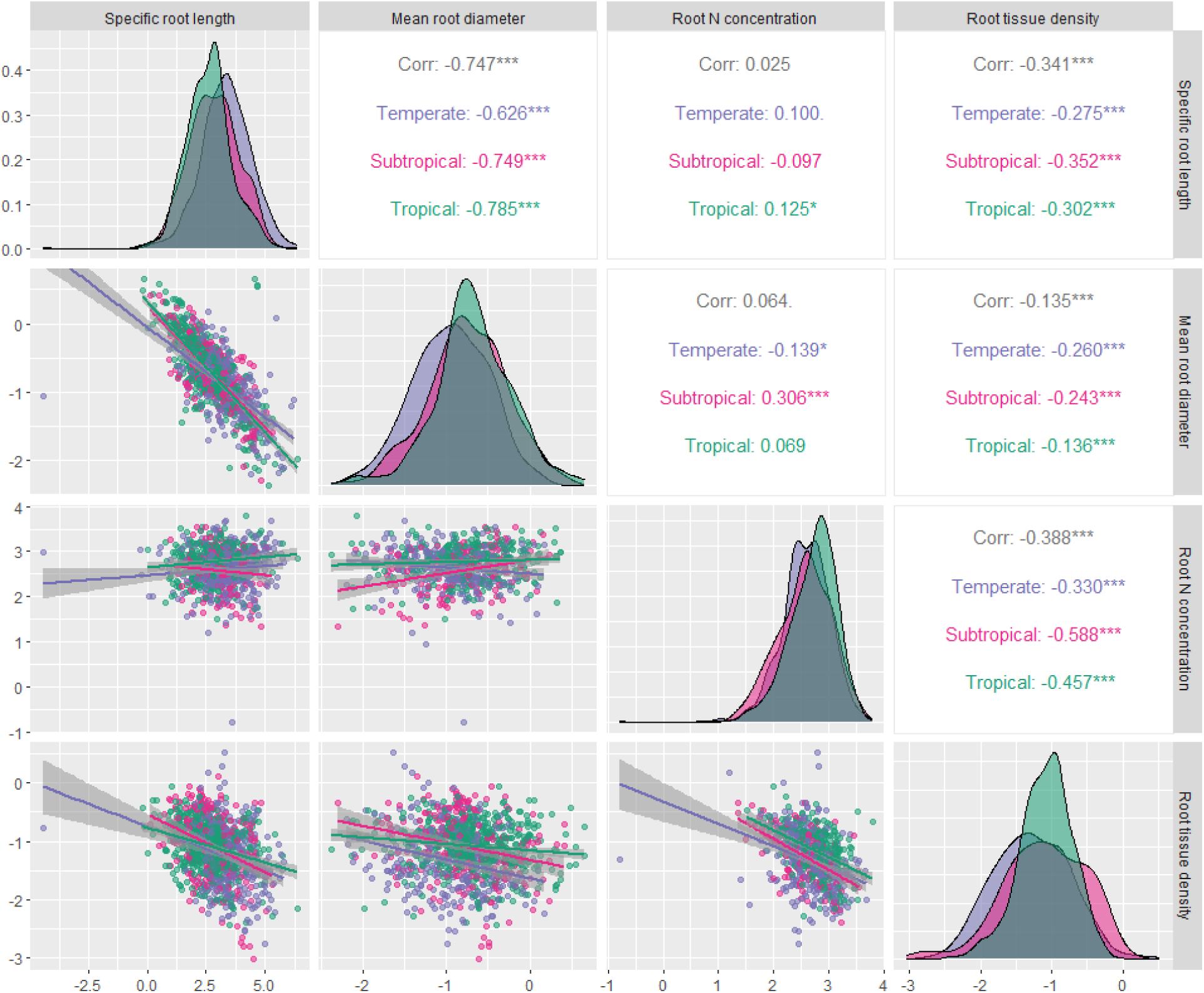
Pearson correlations between root functional traits for woody species.

**Figure S9.**
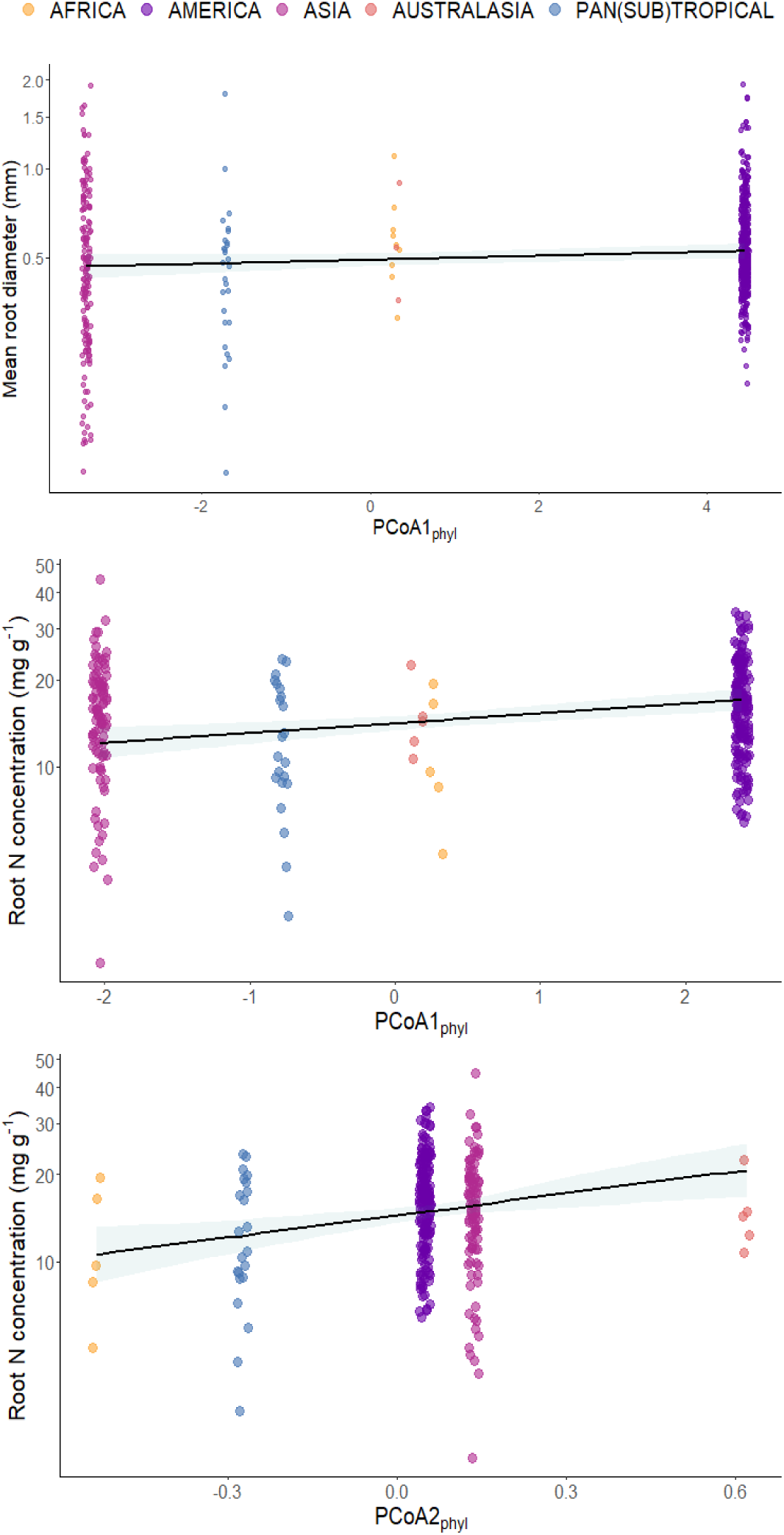
Phylogenetic relatedness explaining root trait in the tropics. The role of phylogenetic relatedness (PCOA1_phyl_ , PCOA2_phyl_), i.e., evolutionary distance between the flora represented in our dataset from different continents, explaining interspecific variation in mean root diameter and root nitrogen (N) concentration. Species from continents that are similar phylogenetically share similar specific root length and root tissue density values.

**Figure S10.**
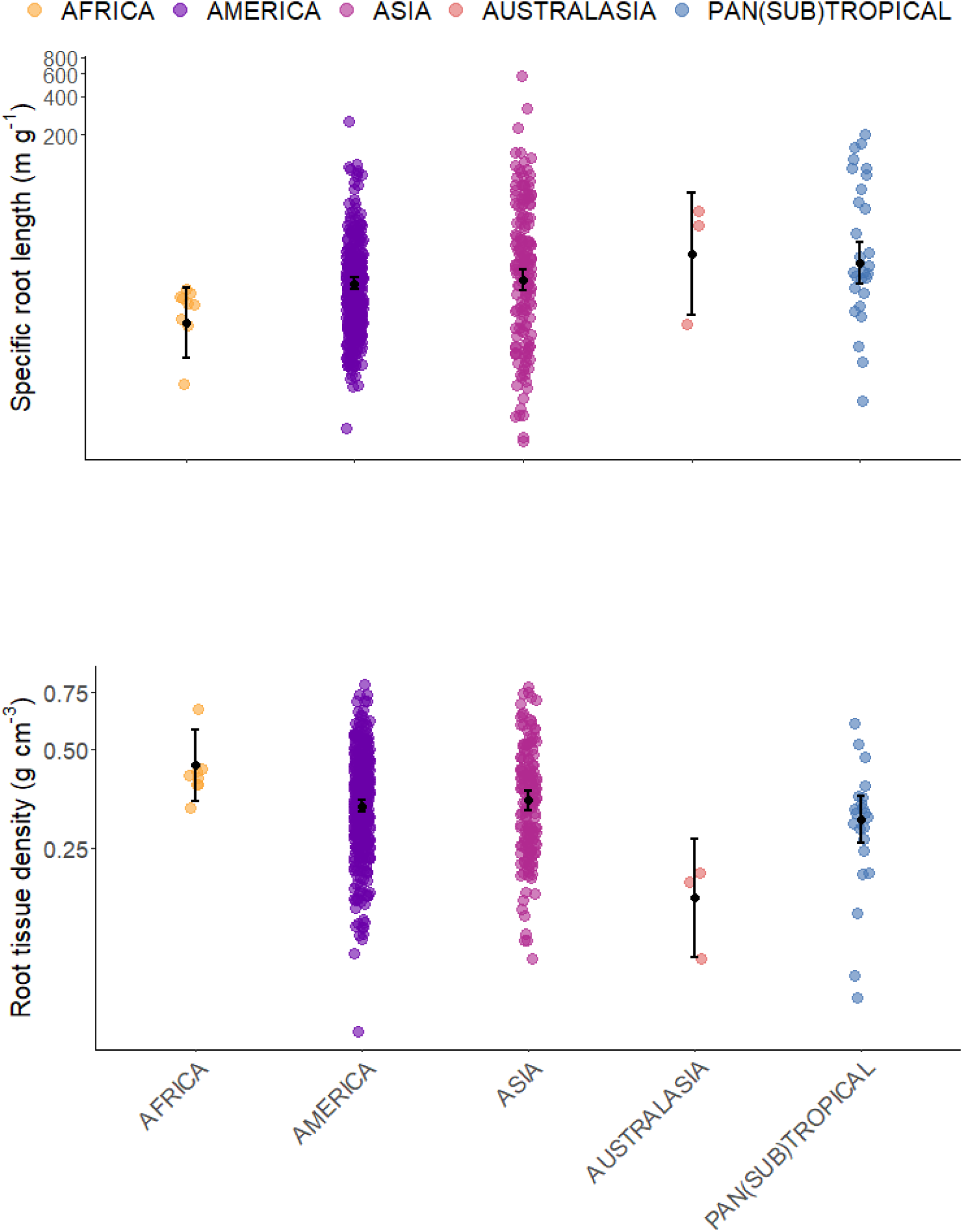
The role of continents explaining root traits in the tropics.

## Supplementary Tables

**Table S1.** Minimum (Min), first and third quartile (1sr Qu. and 3rd Qu., respectively) median, mean, and maximum (Max), values of climatic and soil data used to calculate species’ niches from subtropical and tropical species. In addition, total soil phosphorus (P) data at a 0.5-degree resolution were retrieved from the Global Gridded Soil Phosphorus Distribution Maps (Yang *et al*., 2014), but as these data were unavailable for 20 (sub)tropical species and for 12% of the species occurrence observations (*i.e.,* 44,688) and were correlated with cation exchange capacity (Pearson r = 0.66, p-value < 0.001; Fig S2), we did not include soil P data in our analyses.

| Variables | Min | 1sr Qu. | Median | Mean | 3rd Qu. | Max |
| --- | --- | --- | --- | --- | --- | --- |
| **Climatic variables** |  |  |  |  |  |  |
| Annual mean temperature (°C) | -9.7 | 19.1 | 22.9 | 22.1 | 25.5 | 31.2 |
| Annual precipitation (mm) | 0 | 861 | 1386 | 1578 | 2189 | 10216 |
| Precipitation wettest month (mm) | 0 | 155 | 235 | 241.7 | 336 | 2519 |
| Precipitation driest month (mm) | 0 | 6 | 28 | 44.64 | 60 | 653 |
| Temperature seasonality | 0.68 | 6.8 | 18.3 | 27 | 44.4 | 100.6 |
| Maximum Temperature Warmest month | 35 | 27.6 | 29.9 | 29.8 | 32.0 | 47.6 |
| Min temperature coldest month | -26.8 | 8.0 | 15.3 | 14.1 | 20.5 | 26.7 |
| **Soil variables** |  |  |  |  |  |  |
| Topsoil sand fraction (% wt) | 0.75 | 34.7 | 45.0 | 45.6 | 55.1 | 98.2 |
| Topsoil silt fraction (% wt) | 0 | 22.1 | 27.0 | 27.2 | 32.2 | 65.4 |
| Topsoil clay fraction (% wt) | 0 | 19.6 | 23.6 | 27.0 | 34.0 | 85.0 |
| Topsoil organic carbon (% weight) | 0.09 | 0.86 | 1.17 | 2.25 | 1.85 | 38.17 |
| Topsoil cation exchange capacity (cmol/kg) | 0.5 | 9.2 | 12.0 | 16.2 | 20.0 | 80.0 |
| Total soil phosphorus (g m<sup>-2</sup>) | 45.0 | 230.3 | 321.9 | 404.8 | 485.2 | 1577 |

\*standard deviation $\times 100$ .

**Table S2.** Principal component analyses of root traits using 667 woody species.

|  | PC1 | PC2 | PC3 | PC4 |
| --- | --- | --- | --- | --- |
| Standard deviation | 1.341 | 1.194 | 0.817 | 0.329 |
| Proportion of Variance | 0.450 | 0.356 | 0.167 | 0.027 |
| Cumulative Proportion | 0.450 | 0.806 | 0.973 | 1.000 |
| **Loadings** |  |  |  |  |
| Specific root length | -0.670 | 0.290 | -0.191 | 0.655 |
| Mean root diameter | 0.718 | 0.024 | -0.181 | 0.671 |
| Root tissue density | -0.143 | -0.703 | 0.606 | 0.342 |
| Root nitrogen concentration | 0.118 | 0.648 | 0.750 | 0.052 |

**Table S3.** Principal component analysis of climatic variables for species-specific climatic niches (*position*) for 820 subtropical and tropical species.

|  | **PC1** | **PC2** | **PC3** |
| --- | --- | --- | --- |
| Standard deviation | 2.036 | 1.318 | 0.885 |
| Proportion of Variance | 0.586 | 0.248 | 0.112 |
| Cumulative Proportion | 0.586 | 0.834 | 0.946 |
| **Loadings** |  |  |  |
| Mean annual temperature | 0.365 | **0.496** | -0.134 |
| Mean annual precipitation | **0.456** | -0.123 | 0.363 |
| Precipitation in the wettest month | 0.406 | 0.008 | **0.507** |
| Precipitation in the driest month | **0.378** | -0.354 | 0.220 |
| Temperature seasonality | -0.387 | 0.206 | **0.604** |
| Maximum temperature in the warmest month | 0.017 | **0.738** | 0.168 |
| Minimum temperature in the coldest month | **0.448** | 0.162 | -0.388 |

**Table S4.** Principal component analysis of soil variables for species-specific soil niches (position) for 820 subtropical and tropical species.

|  | PC1 | PC2 | PC3 |
| --- | --- | --- | --- |
| Standard deviation | 1.438 | 1.301 | 0.972 |
| Proportion of Variance | 0.414 | 0.338 | 0.189 |
| Cumulative Proportion | 0.414 | 0.752 | 0.941 |
| **Loadings** |  |  |  |
| Topsoil sand fraction | -0.467 | **0.561** | -0.057 |
| Topsoil silt fraction | 0.539 | 0.019 | **0.626** |
| Topsoil clay fraction | 0.061 | **-0.693** | -0.426 |
| Topsoil organic carbon | 0.385 | 0.369 | **-0.629** |
| Topsoil cation exchange capacity | **0.582** | 0.261 | -0.165 |

**Table S5.**
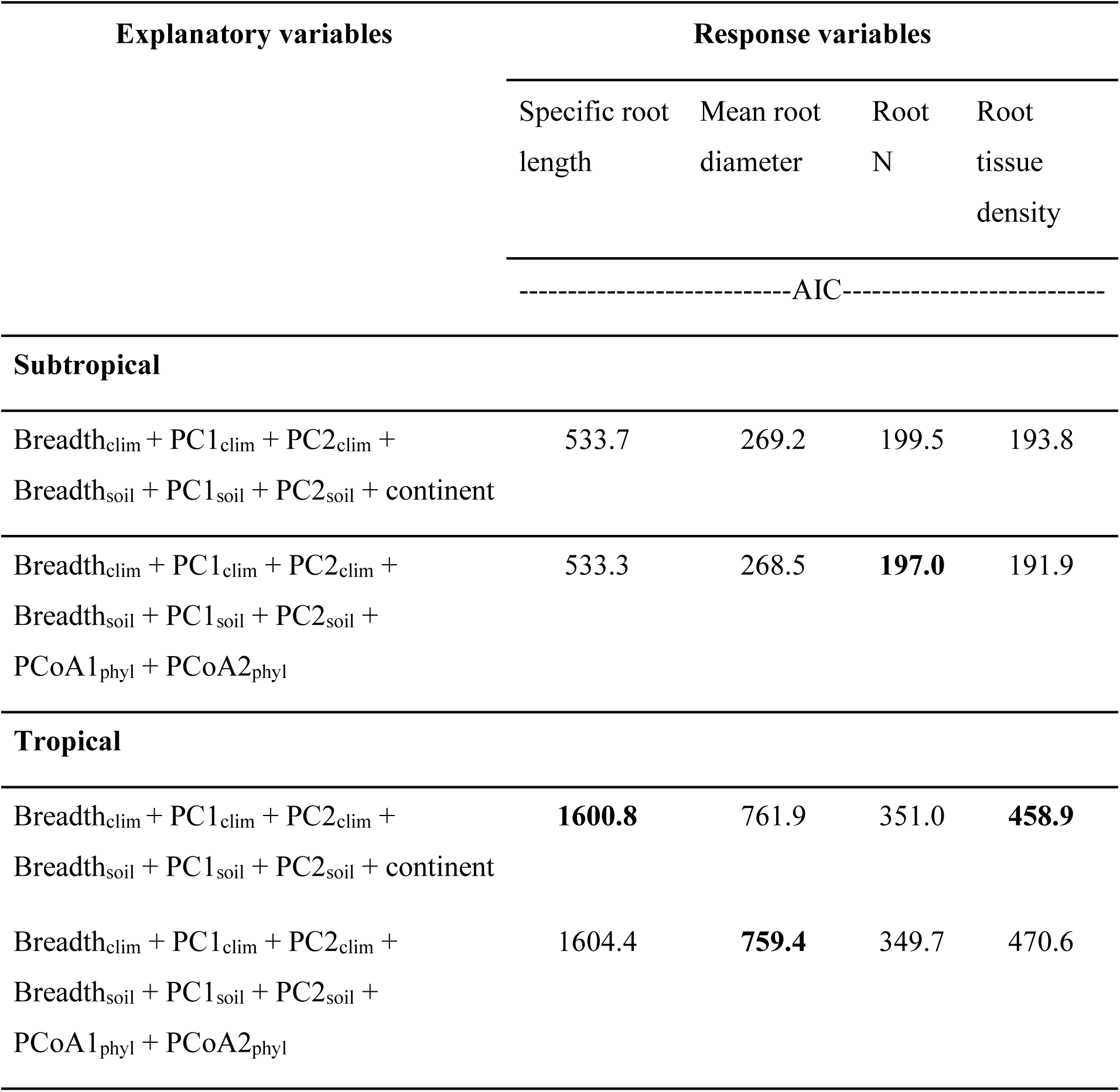
Model selection within biome variation (subtropical and tropical).

| Explanatory variables | Response variables |  |  |  |
| --- | --- | --- | --- | --- |
|  | Specific root length | Mean root diameter | Root N | Root tissue density |
|  | -----AIC----- |  |  |  |
| Subtropical |  |  |  |  |
| Breadth<sub>clim</sub> + PC1<sub>clim</sub> + PC2<sub>clim</sub> +<br>Breadth<sub>soil</sub> + PC1<sub>soil</sub> + PC2<sub>soil</sub> + continent | 533.7 | 269.2 | 199.5 | 193.8 |
| Breadth<sub>clim</sub> + PC1<sub>clim</sub> + PC2<sub>clim</sub> +<br>Breadth<sub>soil</sub> + PC1<sub>soil</sub> + PC2<sub>soil</sub> +<br>PCoA1<sub>phyl</sub> + PCoA2<sub>phyl</sub> | 533.3 | 268.5 | **197.0** | 191.9 |
| Tropical |  |  |  |  |
| Breadth<sub>clim</sub> + PC1<sub>clim</sub> + PC2<sub>clim</sub> +<br>Breadth<sub>soil</sub> + PC1<sub>soil</sub> + PC2<sub>soil</sub> + continent | **1600.8** | 761.9 | 351.0 | **458.9** |
| Breadth<sub>clim</sub> + PC1<sub>clim</sub> + PC2<sub>clim</sub> +<br>Breadth<sub>soil</sub> + PC1<sub>soil</sub> + PC2<sub>soil</sub> +<br>PCoA1<sub>phyl</sub> + PCoA2<sub>phyl</sub> | 1604.4 | **759.4** | 349.7 | 470.6 |

